# OmniPath Metabo: chemical structures, interactions and mechanisms to study the metabolome

**DOI:** 10.64898/2026.06.18.733117

**Authors:** Jonathan Schaul, Yunfan Bai, Daniele Bottazzi, Jannik Franken, Toby Lawrence, Nicolàs Palacio-Escat, Edwin Carreño, Macabe Daley, Lejla Gul, Ardic Sahin, Diego Mañanes, Balazs Bohar, Aurelien Dugourd, Tamas Korcsmaros, Denes Turei, Christina Schmidt, Julio Saez-Rodriguez

**Author notes:** Shared corresponding authors. Shared first authors (Jonathan Schaul, Yunfan Bai, Daniele Bottazzi, Jannik Franken, Toby Lawrence).

## Abstract

Mechanistic and functional analysis of omics data largely relies on the incorporation of prior knowledge; however, connecting metabolomics data and knowledge is a major methodological challenge. This is largely driven by the diverse prior knowledge being fragmented across many databases requiring the merging of different database records across chemical structures, identifiers, and varying levels of structural specificity. Hence, this limits mechanistic interpretation and functional characterisation of the metabolome.

Here, we present OmniPath Metabo, a comprehensive, harmonized, metabolome-centric database covering metabolites, lipids, food-derived compounds, and small molecule drugs, along with their associated receptors, transporters, enzymes, reactions, allosteric regulators, and disease associations. OmniPath Metabo harmonizes attributes using controlled vocabularies and ontologies, structures and built-in cheminformatics to map identifiers and track ambiguity. OmniPath Metabo is built directly from 40+ original resources and is freely accessible via an interactive web app and API at metabo.omnipathdb.org. OmniPath Metabo enables dynamic, context-specific construction of subnetworks to serve dedicated purposes, such as cell-cell communication or integrated multi-omics metabolite-driven regulation, connecting reactions, allosteric regulation, metabolite-receptor and metabolite-transporter interactions. Combining it with the over 170 other resources in OmniPath, it can be used for integrated networks of signaling, gene regulation, and metabolism. We showcase the application of OmniPath Metabo by analysing publicly available metabolomics data of lung cancer cell lines and metabolic footprints to mutational patterns.

In summary, OmniPath Metabo transforms fragmented resources into a harmonised prior knowledge framework for a mechanistic and functional analysis of the metabolome.

## 1. Introduction — Towards a consistent, comprehensive knowledgebase

Mechanistic analysis of omics data largely relies on the incorporation of prior knowledge, yet the integration of metabolism-focused, structured prior knowledge remains a challenge, limiting the mechanistic hypotheses and biological interpretation we are able to extract from metabolomics data. Indeed, compared with transcriptomics and proteomics, metabolomics currently faces three major challenges:

First, the integration of metabolome knowledge, which includes metabolites, lipids, small molecule drugs and food-derived compounds, is complicated by the low adoption of standardized reporting and inconsistent use of metabolite identifiers [1, 2]. While for genes and proteins mapping everything to the same namespace, e.g. primary gene symbols, results in a satisfactory solution for most applications, small molecules cannot be mapped easily by names. Each feature and database record has its own level of structural specificity, from fully defined stereoisomery to varying constitution [1–3]. Since database identifiers point to structures with different levels of specificity, traditional ID translation results in one-to-many and many-to-many mappings, or simply fails to reach the corresponding database record. Database knowledge is essential for virtually all analysis of metabolomics data, hence issues in integrating these two severely impact all downstream analyses. Efforts have been made to improve metabolite cross-database integration, primarily at the level of ID mapping. For example, RaMP-DB 2.0 integrates identifiers across HMDB, ChEBI, PubChem, KEGG, and additional resources while applying molecular-weight-based controls to reduce incorrect mappings [4]. Nevertheless, no current framework comprehensively resolves the issues described here, or provides all required tools and resources for a unified solution.

Second, prior knowledge covering the metabolome remains scattered across numerous databases with distinct scopes, formats, identifier systems, curation strategies, and update frequencies [4]. Consequently, integrating these resources into computational workflows often requires substantial manual effort. Moreover, most resources focus on metabolism, building on well established databases such as KEGG [5] or Reactome [6], with limited integration of receptor-mediated signaling, transporter activity, intercellular communication, allosteric regulation, dietary compounds, microbiome-derived metabolites, or pharmacological interactions. This represents a major limitation because metabolite functionality stretches far beyond classical metabolic intermediates: they can act as signaling molecules, receptor ligands, transporter substrates, regulatory cofactors, drugs, and mediators of tissue and cell-cell communication [7–9]. As metabolomics becomes increasingly integrated with other omics technologies, there is a growing need for metabolite-centered prior knowledge capable of linking metabolic states to signaling, transcriptional regulation, and disease phenotypes.

Third, conventional metabolomics interpretation strategies remain largely constrained by the pathway-centric view [7], thus portraying metabolism only as a passive bystander of gene regulation. Moreover, metabolic pathways are often linear rather than co-regulated, metabolite abundances reflect both production and consumption fluxes, and many metabolites are rapidly transported between tissues, extracellular compartments, and biofluids [7]. Recently, it has become clear that the metabolome interacts with and modulates the cellular phenotype through regulation of other omics layers (e.g. epigenome, transcriptome, proteome) via for example signalling [10–14]. Enabling the analysis of the metabolome as an active regulator is limited by incomplete database knowledge of pathways, tissue and compartment specificities, and non-canonical metabolite functions such as receptor signaling, transporter-mediated communication, and allosteric regulation.

Together, these limitations of (**i**) the unresolved problems of metabolite identifier mapping, (**ii**) fragmented and poorly interoperable prior knowledge resources, and (**iii**) the conceptual limitations of pathway-centric analysis frameworks, highlight the need for integrative metabolite-centric prior knowledge frameworks capable of supporting mechanistic and context-aware analyses beyond canonical pathway enrichment.

Here, we present **OmniPath Metabo** (metabo.omnipathdb.org), a comprehensive metabolite-centric prior knowledge resource designed to enable mechanistic biological interpretation focused on the metabolome. This major extension of the originally gene focused OmniPath prior knowledge framework [15, 16] enables the harmonized integration of heterogeneous metabolite-centric resources. OmniPath Metabo integrates metabolites, lipids, food-derived compounds, drugs, ligands, receptors, transporters, enzymes, reactions, allosteric regulators, and disease associations into a unified database. It is built directly from original databases through a modular integration pipeline using controlled vocabularies, ontologies, and chemical structure harmonization. Importantly, OmniPath Metabo systematically resolves cross-database identifiers while explicitly tracking structural ambiguity and uncertainty, enabling transparent and reproducible metabolite mapping across resources. In addition, identifiers and names are comprehensively linked to chemical structures and cheminformatics functionalities through RDKit [17], supporting structure-aware analyses and flexible metabolite matching strategies. OmniPath Metabo also enables the generation of dynamic and context-specific subsets, to serve dedicated purposes, such as intercellular cell-cell communication, combined signaling-metabolism networks or integrated multiomics metabolite-driven regulation. Indeed, this has enabled the dynamic extension of e.g. intercellular metabolite-protein interactions and multi-layer network of metabolite-driven regulation for metabolite-mediated signaling and cell-cell communication. Together, OmniPath Metabo merges fragmented metabolic resources into a unified prior knowledge framework for mechanistic and systems-level analysis of metabolism within multi-omics contexts.

## 2. Results

### 2.1. OmniPath Metabo integrates metabolome and systems biology resources through a modular harmonization pipeline

To address the fragmentation of metabolite-centered prior knowledge, we developed a modular data integration framework that systematically combines 40+ resources into a unified database architecture (Figure 1). These include databases of small molecules and lipids, metabolic reactions and pathways, metabolite-protein interactions, receptor and transporter biology, disease associations, food-derived compounds, protein function, molecular complexes, and biological ontologies (Figure 1a; Table 1; Supplementary Table 1).

**Figure 1.**
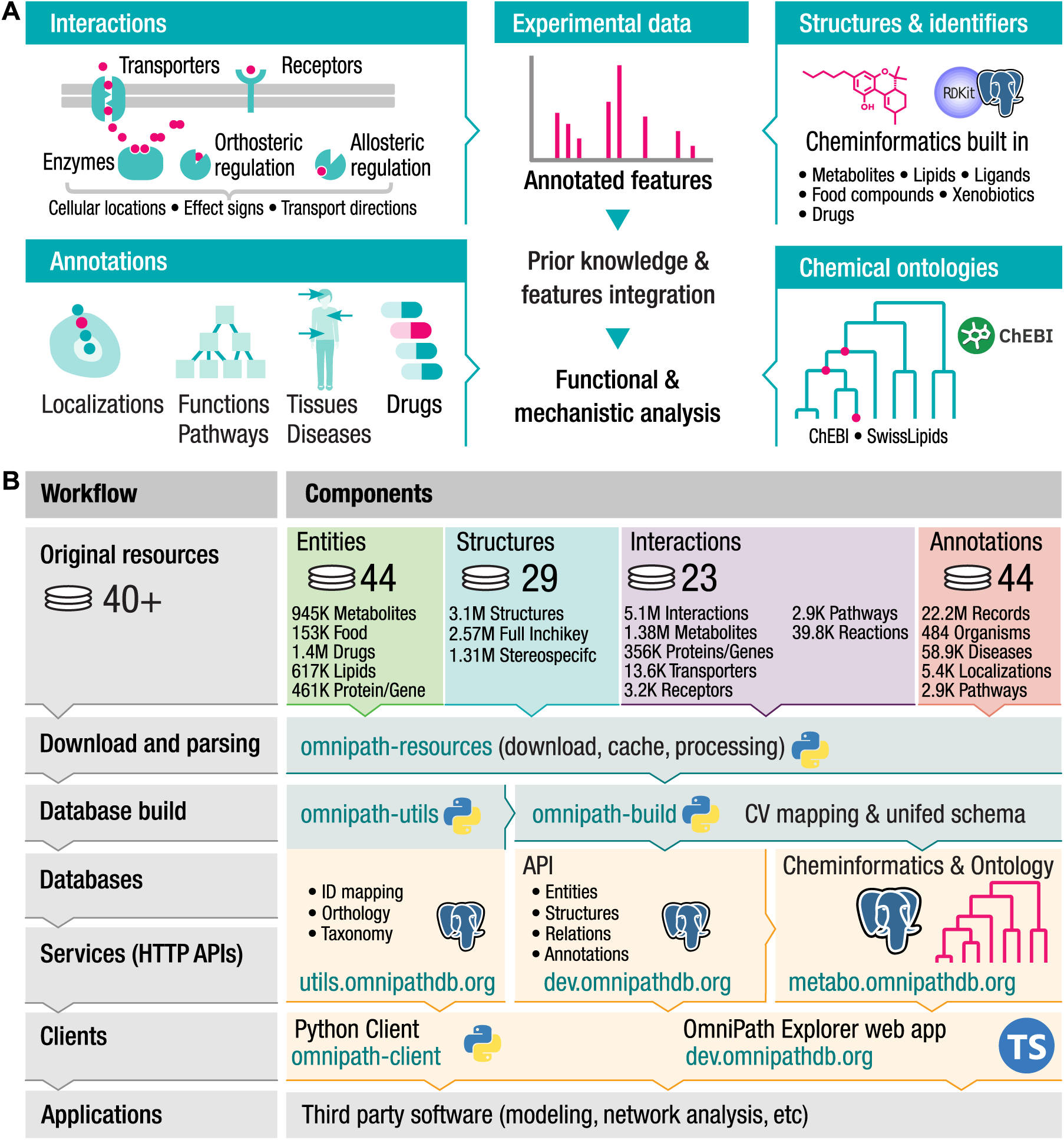
OmniPath Metabo database workflow and architecture. **A**) High level overview of Omnipath Metabo. The database combines various mechanistic interac-tions, annotations, chemical structures and ontologies. **B**) The left side shows the stages of the prior knowledge processing workflow. The right side shows the major software components and their roles in the workflow. Resource specific parsing code and direct access to original resources is implemented in omnipath-resources; using these parsers, omnipath-build and omnipath-utils build the OmniPath and OmniPath Utils databases, respectively. The omnipath-metabo package adds cheminformatics and chemical ontologies to the main OmniPath database. The database contents are available by three public web APIs: Utils, main, and Metabo, all accessible from Python by the omnipath-client package. The OmniPath database is browseable by an interactive web app.

**Table 1.**
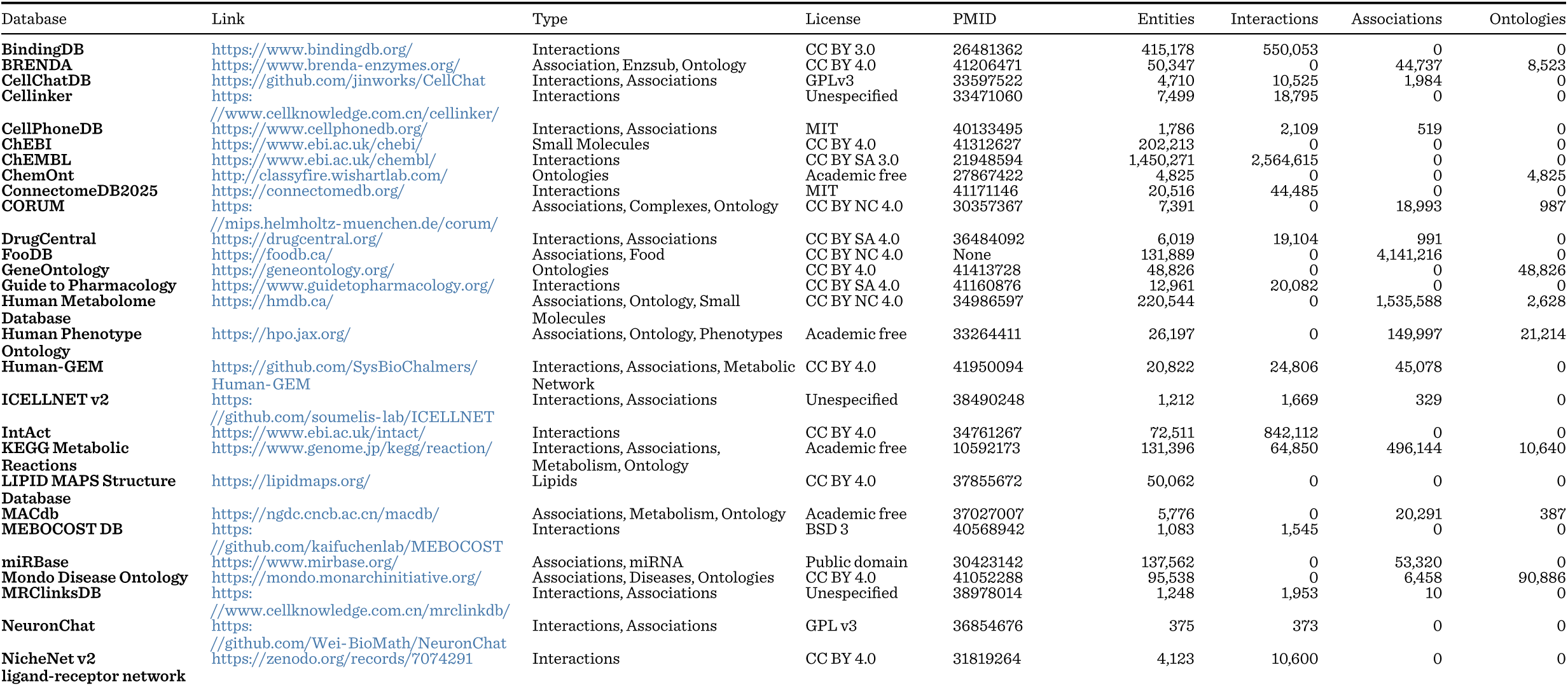
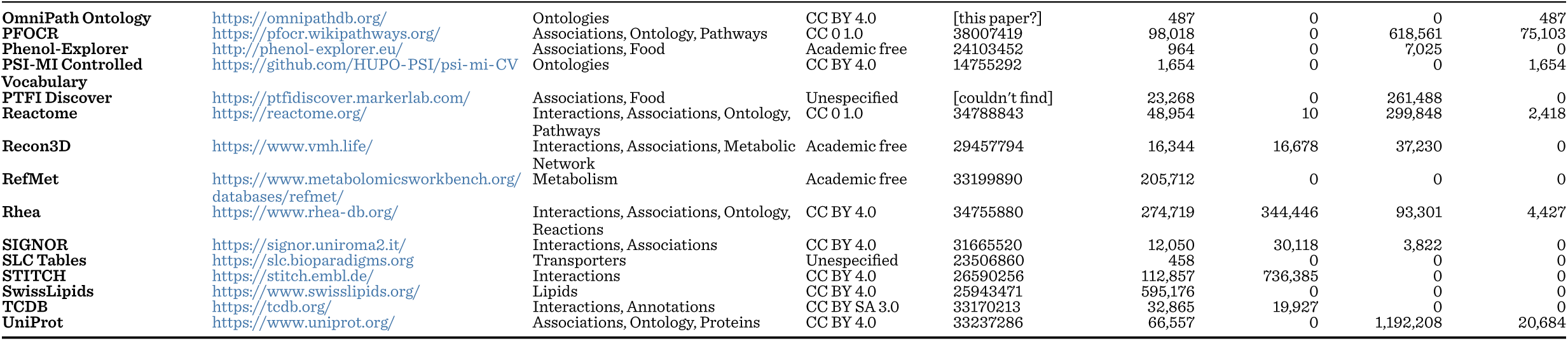
List of main resources implemented in OmniPath Metabo.

The build pipeline separates resource ingestion, normalization, entity resolution, and graph generation into independent stages (Figure 1b). Individual source databases are first processed through resource-specific input modules that convert heterogeneous source formats into a common internal format. This normalization enables the representation of fundamentally different biological entities, such as metabolites, proteins, chemicals, reactions, diseases, etc. in a common, ontology-backed schema, integrating them into a unified graph. Importantly, all records retain complete source provenance throughout the integration process, ensuring traceability from the final database back to the original record. A key feature of the design is the explicit separation between source evidence and canonical graph records. Source-derived observations are first stored as resource-specific evidence objects before being resolved into shared entities and relationships (Figure 1b). This strategy allows records originating from independent databases to be merged when they refer to the same biological entity, while preserving all supporting evidence and provenance information (details in methods section **4.2.5**). Consequently, users can simultaneously benefit from increased coverage through integration and from transparent access to the original supporting resources.

For resolution of chemical entities, we always rely on InChI keys when available; entities that cannot be represented as InChI are canonicalized by nodes of the ChEBI ontology [18, 19]; databases that do not provide structures are unambiguously translated to InChI keys or ChEBI whenever possible; and, for lipids, we use Goslin [20] to normalize names and traverse levels of structural specificity. We use RDKit [21] and Goslin to classify structures by levels of structural specificity. Having RDKit cheminformatics, the ChEBI ontology, and the Goslin lipid parser integrated in our database not only helps build OmniPath Metabo, but also provides a powerful toolkit to address difficult methodological problems in computational work with chemicals.

OmniPath Metabocontentisbrowseableinawebapp(https://dev.omnipathdb.org), which supports interactive exploration of molecular entities, relations, and ontologies (Figure 2, Supplementary Figure S1). Through the Explore and Selection views, users can search and filter entities or relations, inspect supporting details and evidence, and iteratively refine selected subsets using shared ontology terms. By linking chemicals, proteins, pathways, ontology terms, and their relations in a single workflow, the interface enables exploratory analysis, hypothesis generation, and biological interpretation of results.

**Figure 2.**
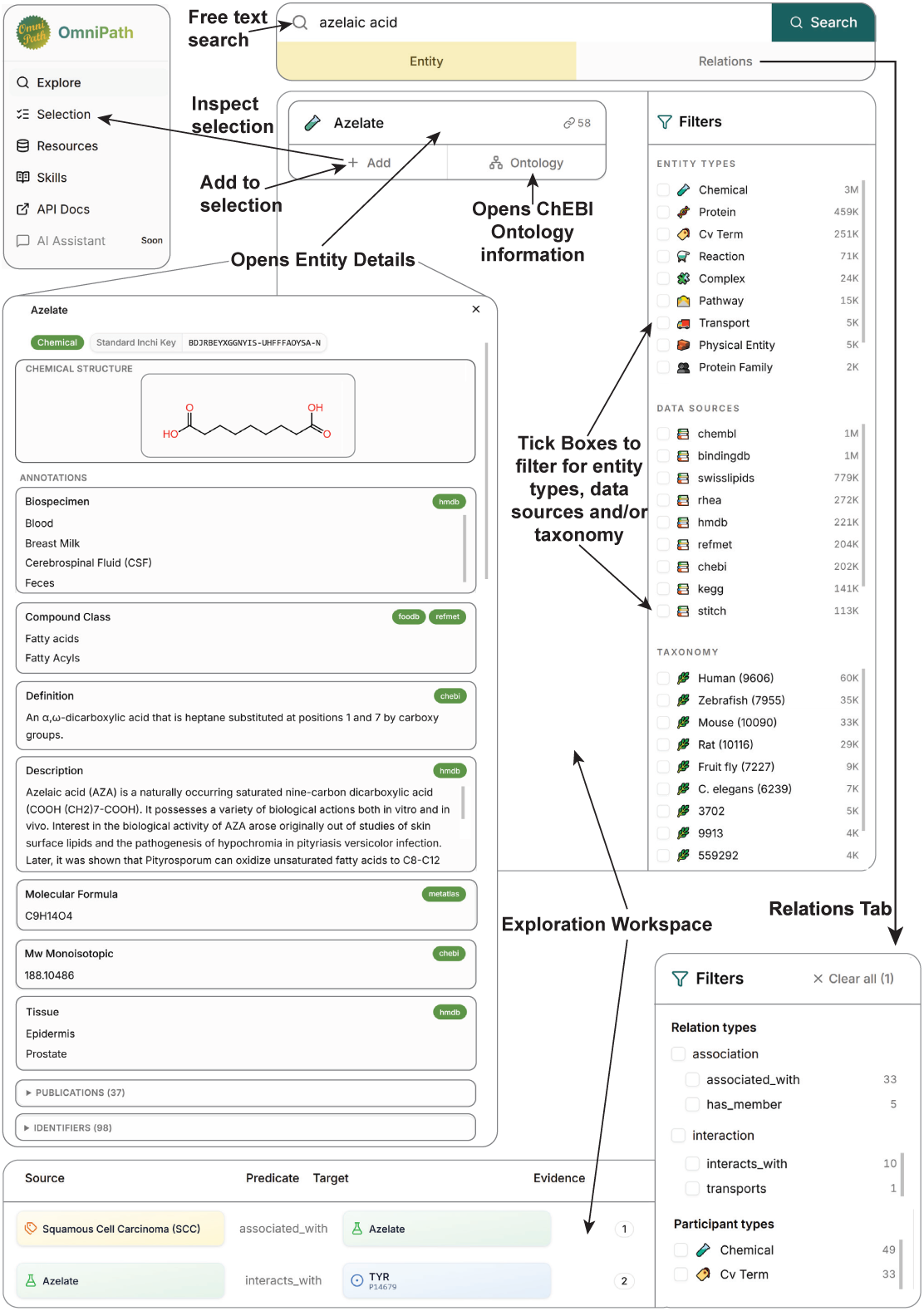
OmniPath Metabo web interface. Screenshots illustrating key components of the OmniPath Metabo web application. The interface combines entity-centric, relation-centric, and ontology-based exploration within a unified workflow. **Top left**: persistentnavigation sidebar providing access to the Explore, Selection and Resources views. **Top right**: search interface with faceted filters for querying entities and relations. **Middle left (“Entity Details”)**: detailed view of a selected entity (azelaic acid shown as an example), including identifiers, annotations, cross-references, and chemical structure visualization. **Bottom (“Relations Tab”)**: exploration of biological relations associated with the queried entity, including source and target entities, interaction types, and supporting evidence. **Bottom right (“Filters of Relations Tab”)**: filtering options enabling restriction of results by entity type, taxonomy, relation type, and source database. Together, these components support interactive exploration of metabolites, proteins, pathways, ontology terms, and their associated biological relationships.

### 2.2. OmniPath Metabo integrates information from diverse resources

The current OmniPath Metabo release integrates information from 40+ resources, representing over 4 million entities (e.g. lipids, metabolites, food compounds, drugs proteins, genes etc.), almost 3 million unique structures (InchiKeys) connected to 16 million identifiers, almost 15 million interactions (e.g. metabolite-protein, metabolite-pathway), and over 800 thousand associations (e.g. between complex and its members) (Figure 3a; Supplementary Table 2). The most numerous types of entities are lipids, food compounds, and drugs, (Figure 3a), the largest contributing resources being ChEMBL [22], SwissLipids [23] and BindingDB [24] (Supplementary Figure S2, left top panel). OmniPath Metabo also incorporates literature references including DOI, PMC and PubMed IDs with over 200 thousand (Figure 3a) and the majority originating from ChEMBL [22], Pathway Figure OCR (PFOCR) [25] and BindingDB [24] (Supplementary Figure S2, right panel). Chemical entities can be separated into structurally defined compounds with standard InChI keys and entities without structural information (Figure 3b). Based on their InChI keys, compounds can be further stratified by their levels of structural specificity. In Figure 3b, we show the count of chemicals with stereospecific cis/trans specific, single constitution without stereo information and variable constitution (e.g. structure with a variable moiety) entities. The structures play a central role within the entity resolution performed within OmniPath Metabo, which uses dedicated identifier resolver tables constructed from authoritative source-specific mappings (Methods 4.2.4). With this OmniPath Metabo overcomes a major obstacle in metabolism-related database integration, the lack of consistent metabolite identifiers across resources [1, 2]. For metabolites and lipids, identifiers from resources including ChEBI [18], HMDB [26, 27], PubChem [28], LipidMaps [29, 30], SwissLipids [23], RefMet [31], and others are resolved to canonical chemical representations whenever possible. This refers to structurally defined compounds where standard InChIKeys serve as the primary integration layer, enabling records originating from multiple databases to converge on a common chemical representation. When complete structural information is unavailable, additional curated cross-reference mappings are used while preserving ambiguity information (Methods 4.2.4). For gene products including proteins, identifiers from multiple namespaces are unified to NCBI Entrez Gene IDs [32]. Across all incorporated resources, the resolver framework contains over 3 million unambiguous chemical mappings, providing a scalable foundation for cross-resource metabolite integration. Together this leads to a highly connected network between the original resources (Figure 3c). The interactions within OmniPath Metabo span signalling, gene regulation, transport, metabolism, and various regulatory interactions of chemicals (Figure 3d). Unlike entities, which show extensive cross-resource overlap (Figure 3c), interactions are more specific for the source, with many relations contributed by only one resource (Figure 3e).

**Figure 3.**
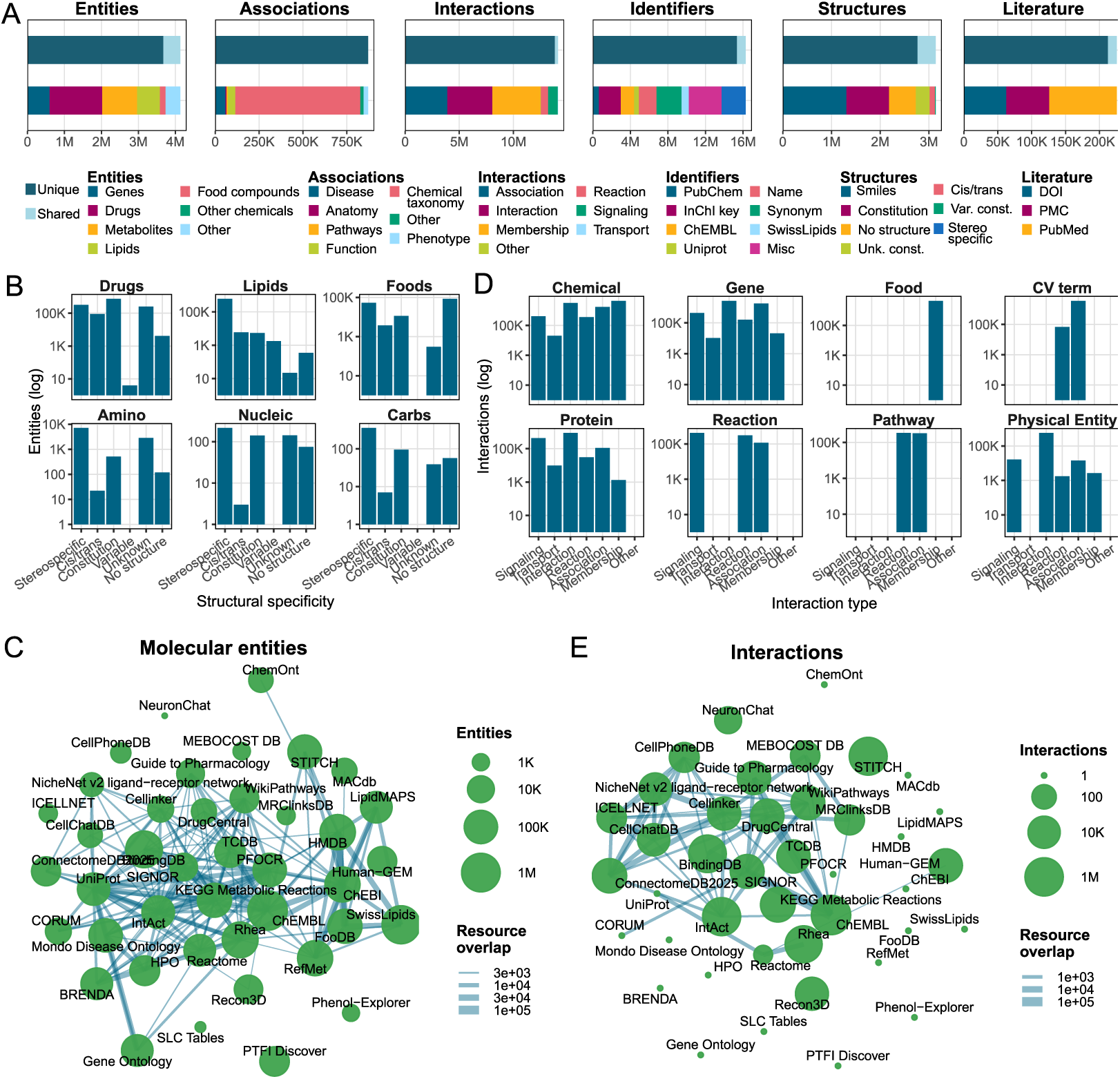
OmniPath Metabo database content. **A**) Number of records by type (entities, associations, interactions, identifiers, structures and literature references). Colors of the top bars show the unique (dark; occurs only in a single resource) and shared (light; in multiple resources) records, while colors of the lower bars show major categories within each record type. **B**) Resource sizes and overlaps: node sizes are proportional to the number of entities (left) and interactions (right), while edge widths are proportional to the pairwise overlap of resources. **C**) Level of structural specificity of chemical entities. Number of entities (log) in each specificity level by major classes of chemicals. “Unknown” are unknown constitutions whilst “No Structure” did not provide structural information. **D**) Interaction types by entity types: number of interactions (log) by interaction types for major classes of entities.

To sum up, OmniPath Metabo includes a wealth of metabolism-related knowledge and its modular architecture enables new resources to be incorporated without redesigning the underlying schema, facilitating continuous expansion of the database as additional metabolomics knowledge becomes available.

### 2.3. OmniPath Metabo derives application-specific subsets to study to study multi-omics metabolite-driven regulation accessible via API

The contents of OmniPath Metabo is largely unfiltered, harmonized, and stored in a simple general schema, enabling the compilation of application-specific prior knowledge. Currently we provide one such datasets through the API.

Applying this, we implement a network of metabolism regulation, as one of the major challenges in multi-omics integration is the limited representation of metabolism within mechanistic prior knowledge networks. Metabolism is typically reduced to canonical biochemical reactions connecting metabolites and enzymes. As a consequence, metabolites are often treated as passive downstream readouts of pathway activity rather than active regulators of cellular signaling and communication. Hence, we have built a prior knowledge network to study multi-omics metabolite-driven regulation by connecting signaling, transport, enzymatic, regulatory, and biochemical interaction knowledge (Figure 4a; Table 3), basing this on our original concepts presented in our COSMOS framework [33, 34]. Because metabolite-receptor, metabolite-transporter, allosteric regulation, metabolic reaction, subcellular localization, and signaling interactions are integrated into the same interoperable back-end, it makes it possible to dynamically construct causal prior knowledge networks in which metabolites function as active mechanistic entities rather than solely pathway intermediates (Figure 4a). Here, receptor-mediated and transporter-mediated interactions establish direct mechanistic links between extracellular metabolites and intracellular signaling cascades, thereby bridging metabolism with canonical signaling networks. In parallel, the integration of allosteric regulation introduces regulatory feedback mechanisms that are absent from conventional genome-scale metabolic representations, while organelle and compartment annotations provide additional biological context for constraining mechanistic hypotheses.

**Figure 4.**
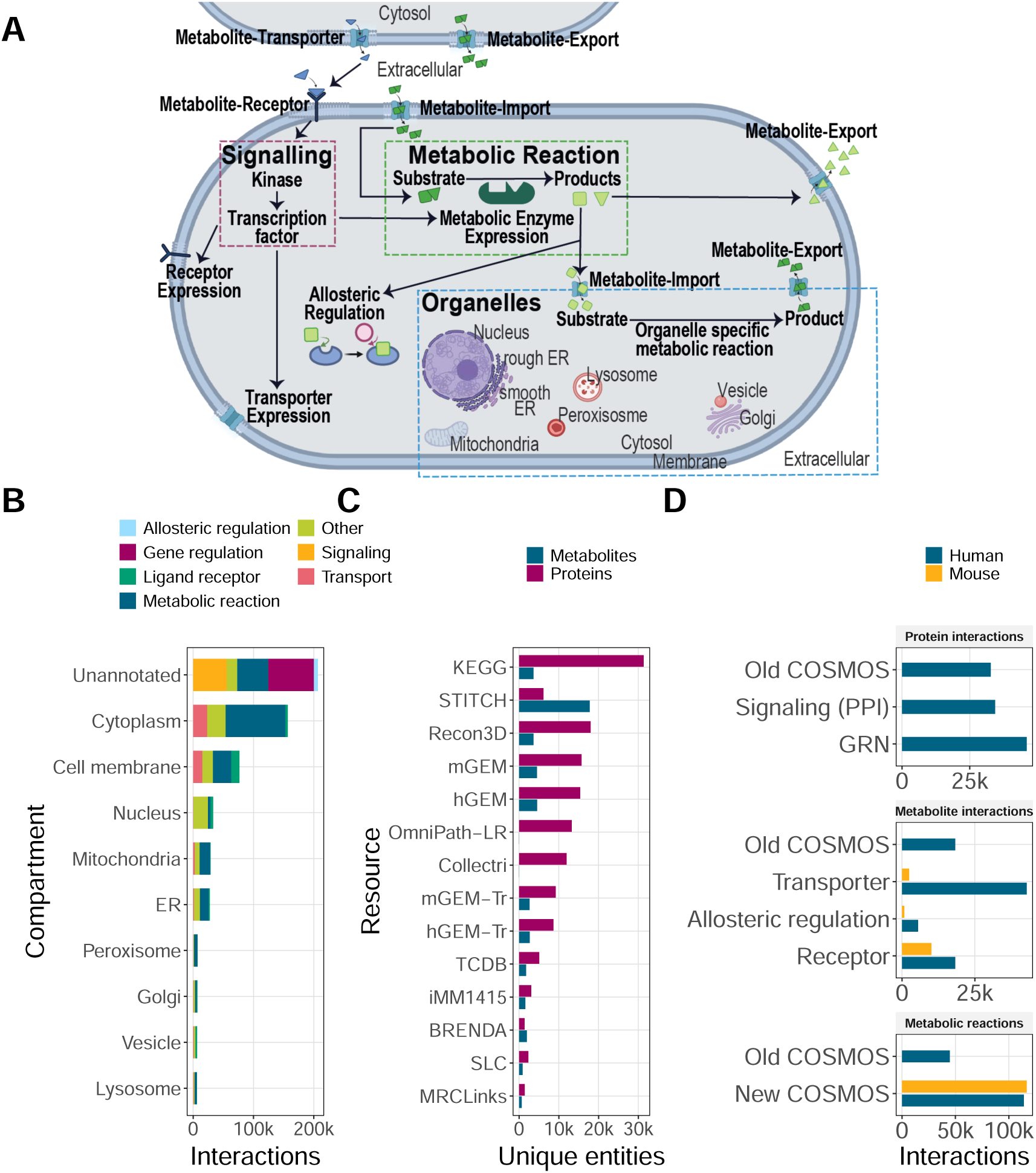
Metabolite mediated interactions in an integrated network of molecular regulation. **A**) Metabolic centric biological entities included with cell-cell communication and intracellular metabolite-driven signalling, allosteric regulation and metabolic reactions. Metabolite import and export as well as metabolic reactions are linked to different cellular organelles. The prior knowledge network to study multi-omics metabolite-driven regulation (COSMOS PKN) includes: **B**) Cellular compartments of interactions. ER: endoplasmic reticulum. **C**) Molecular entities in the COSMOS PKN per data resource. LR: ligand-receptor; Tr: transport. **D**) Comparison of the original COSMOS prior knowledge network (“Old COSMOS”) with the new one (“New COS-MOS”) by interaction type. In the old version, interaction types cannot be distinguished, hence direct comparison of protein and metabolite interactions is not possible (top two facets). GRN: gene regulatory network; PPI: protein-protein interactions.

**Table 3.** Data resources of multi-omics metabolite-driven regulation.

|  | Data Sources |
| --- | --- |
| Transporter | TCDB, SLCtable, Metabolic Atlas, BiGG, MRCLinksdb |
| Receptor | MRCLinksdb, Metabolic Atlas, STITCH |
| Allosteric regulation | BRENDA |
| Metabolite-protein interaction | STITCH |
| Enzyme-Metabolite | Metabolic Atlas, KEGG, BiGG |
| Signaling Network | OmniPath |

The new multi-omics metabolite-driven regulation knowledge includes compartment specific knowledge (Figure 4b) from different sources (Figure 4c). This includes metabolite-mediated interactions between metabolites and proteins (Figure 4c), which can be accessed in a compartmentalized manner. A substantial proportion of interactions that are not annotated to a specific compartment (Figure 4b, unannotated). In total this covers over 200 thousand interactions across nice cellular compartments (Figure 4b). Compared to the original COSMOS prior knowledge network [34], this has substantially expanded the size and coverage of the integrated network covering multi-omics metabolite-driven regulation (Figure 4d). In particular, the inclusion of transporter-metabolite and receptor-metabolite interactions increased the number of metabolite-protein regulatory edges (Figure 4d, Metabolite interactions) introducing previously unrepresented connections between extracellular metabolism and intracellular signaling pathways.

Overall, this provides a more biologically realistic representation of cellular regulation, where metabolism and signaling are modeled as deeply interconnected systems rather than separate functional layers. We anticipate that this framework will be particularly relevant for applications involving cancer metabolism, immunometabolism, microbiome-host interactions, and spatial omics, where metabolites frequently mediate communication between cells and tissues.

### 2.4. Case-study of context-aware interpretation of lung cancer metabolomics using OmniPath Metabo

To demonstrate the utility of OmniPath Metabo for mechanistic interpretation of metabolomics data, we reanalyzed a previously published cancer cell line metabolomics dataset from Shorthouse et al. [35] restricted to lung and pleura-derived cell lines, which represented the largest subgroup in the dataset (∼45% of all cell lines). Because oncogenic drivers frequently co-occur with additional mutations that independently reshape cellular metabolism, we designed the analysis to capture both broad mutation-associated metabolic signatures and more specific driver-associated effects after controlling for major co-mutation patterns by filtering the cell lines for *TP53*-mutant, *CDKN2A*-wild-type, *STK11*-wild-type and (**i**) *KRAS*-wild-type or (**ii**) EGFR-wild-type. For the comparison of *KRAS*-mutant versus *KRAS*-wild-type we filtered for *EGFR*-wild-type cells, whilst for the comparison of *EGFR*-mutant versus *EGFR-*wild-type cell lines we filtered for *KRAS*-wild-type. Differential metabolite abundance analysis was performed and metabolites were ranked by moderated t-statistics to prioritize the most consistently altered features across contrasts (**Supplementary Table 4**). We then used OmniPath Metabo to perform multi-layered contextual interpretation of the resulting metabolite signatures.

First, we explored whether the integrated search and annotation functionalities could rapidly recover biologically and clinically relevant knowledge for individual metabolites. For example, azelaic acid emergedamong the stronglydifferential metabolites, suggesting decreased abundance in KRAS-mutant cell lines but showing increased abundance in EGFR-mutant backgrounds (Figure 5a-b). Through the OmniPath Metabo website search (Figure 3) we learned that azelaic acid (or synonym azelate) is a naturally occurring saturated nine-carbon dicarboxylic acid and occurs in different biospecimens such as blood, sweat and urine, supporting its potential utility as a circulating biomarker. We further found that azelaic acid also naturally occurs in potatoes, asparagus and common beans and can be produced by lipid oxidation (**Supplementary Table 5A-C**). Via the OmniPath Metabo FastAPI, we were also able to extract relations for azelaic acid finding 9 interactions from ChEMBL, Stitch and Drug-Central (Figure 5c) including *TYR* (Tyrosinase) (**Supplementary Table 5D**). However, no interaction we found could explain our observations regarding the different azelaic acid changes in EGFR-mutant and KRAS-mutant cells (Figure 5a-b). Additionally, most associations were cancer associations reported by MACdb [36]. Hence we next used OmniPath Metabo to extract cancer associations reported in connection with azelaic acid and found 17 associations with lung cancer across different sample types (Figure 5d). For the studies that reported those associations, we used the pubmed IDs provided by OmniPath Metabo (**Supplementary Table 5E**) to manually review each study and found no mutation information in the patient cohorts. Given azelaic acid is generated through oxidative cleavage of unsaturated fatty acids [37], its abundance may reflect cellular lipid oxidation processes and be influenced by lipid peroxidation, Reactive Oxygen Species (ROS) handling, fatty acid composition and subsequent metabolic turnover [38]. Oncogenic KRAS has been shown to activate antioxidantprograms thatenhance ROS detoxification in lung cancer cells [39], hence the decrease in azelaic acid could therefore reflect lower accumulation of oxidation products rather than lower production.

**Figure 5.**
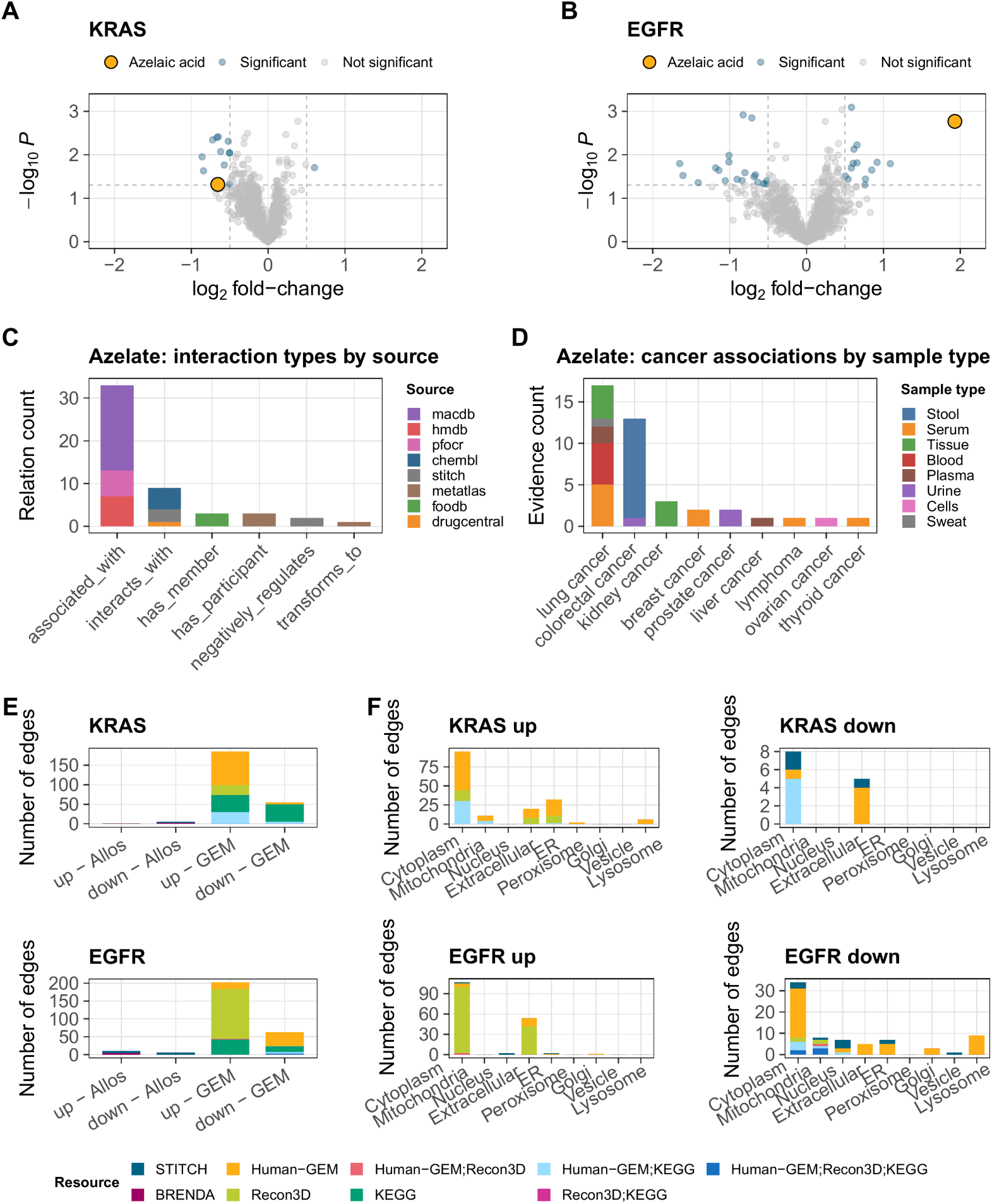
Case-study showcasing how OmniPath Metabo enables investigation of bi-ologically relevant metabolites in cancer cell line metabolomics. **A-B**) Volcano plots showing results of differential metabolite analyses of Shorthouse et al. [35] lung cancer cell lines for TP53-mutant, CDKN2A-wild-type, STK11-wild-type and **A**) EGFR-wild-type cell lines, comparing *KRAS*-mutant versus *KRAS*-wild-type cell lines, and **B**) *KRAS*-wild-type cell lines, comparing *EGFR*-mutant versus *EGFR*-wild-type cell lines. Limma-based differential analysis was performed on log2-transformed intensity data, and volcano plots show log2FC versus -log10(pValue), while highlighting in blue significant observations (pValue <= 0.05). **C-D**) Azelaic acid search results from the Omnipath Metabo FastAPI. **E**) GEM using BiGG, Metabolic Atlas, and KEGG with connections to allosteric enzyme regulation from BRENDA. **F**) Subcellular compartment distribution of PK interactions for differenttially abundant metabolites.

To further explore interactions involving the top altered differential metabolites in the KRAS and EGFR contrasts, we used OmniPath Metabo to integrate metabolite reaction resources from BiGG [40], Metabolic Atlas [41], and KEGG [5], together with knowledge of allosteric enzyme regulation from BRENDA [42] (Figure 5e). We identified connections between differential metabolites and allosteric regulatory interactions and extracted approximately 250 genome-scale metabolic network edges for each contrast. To investigate whether the differentially abundant metabolites exhibited common subcellular patterns, we next examined their compartmental distribution (Figure 5f). Although most metabolites were broadly assigned to cytoplasmic and extracellular compartments, the analysis also revealed more specific associations with organelles such as the endoplasmic reticulum (ER) and lysosome (Figure 5f). By integrating these resources, we generated a metabolism-centred network that captures both metabolic reactions and regulatory interactions. This framework provides an up-to-date and customizable resource for studying metabolite-mediated cell-cell communication and exploring the functional context of differential metabolites.

Taken together, these analyses illustrate how OmniPath Metabo enables the systematic contextualization of metabolomics findings across multiple layers of biological knowledge. Starting from differential metabolite abundance signatures, we were able to rapidly retrieve metabolite-specific annotations, explore molecular and disease associations, integrate metabolic reactions and regulatory interactions, and place metabolites within their subcellular context. The azelaic acid example further demonstrates how OmniPath Metabo can support the generation of mechanistic hypotheses by linking experimental observations to existing knowledge. By interconnecting diverse resources through a unified framework, OmniPath Metabo facilitates biologically interpretable networks, enabling deeper exploration of metabolic phenotypes and the formulation of testable hypotheses from complex metabolomics datasets.

## 3. Discussion

In this work, we present OmniPath Metabo as a metabolite-centric prior knowledge framework that addresses three major limitations in current metabolomics analysis: fragmented biological resources, metabolite identifier mapping, and the conceptual constraints of pathway-centric interpretation. By integrating heterogeneous metabolism, lipids, signaling, transport, regulation, and disease resources into a unified ontology-backed schema, OmniPath Metabo enables context-aware and mechanistic interpretation of metabolomics data within both single- and multi-omics settings.

A key aspect of OmniPath Metabo is the explicit use of chemical structure information as the primary integration layer for metabolite harmonization. In contrast to identifier-centric integration strategies such as RaMP-DB [4], which uses an identifier integration approach, OmniPath Metabo resolves records through standardized structural representations whenever possible. This distinction is important because metabolite identifiers frequently represent compounds at different levels of structural specificity, including stereoisomers, constitutional isomers, or broader compound classes [43]. Identifier-based collapsing can therefore merge biologically distinct entities or propagate incorrect cross-references between databases. To mitigate this, RaMP-DB applies additional safeguards: candidate merges are flagged when molecular weights differ by 10% or more, and known problematic mappings are excluded using manually curated exclusion lists [4]. The current RaMP-DB build contains 44,899 ambiguous mappings, where a RaMP ID is associated with more than one distinct molecular structure. These mappings involve 22,416 unique RaMP IDs (Supplementary Table 3). OmniPath Metabo instead treats structural identity as the canonical integration layer, while explicitly preserving ambiguity when structural resolution is incomplete. This allows users to distinguish between unambiguous structure-level mappings and lower-confidence relationships derived from incomplete annotations, thereby improving transparency and reproducibility in metabolite integration workflows.

The incorporation of an internal RDKit [21]-based structural processor is central to this strategy, just like the integration of the ChEBI ontology [18, 19] and the Goslin [20] lipid name processor. The combination of these tools enables connecting features and records at different levels of structural resolution, while the rich annotations in the database enable doing this in a biological context-aware way. Beyond enabling canonicalization and standardization of molecular structures, the structural processor provides a foundation for future structure-aware analyses directly within OmniPath Metabo. These include similarity searches, substructure matching, lipid class inference, ontology traversal, and structural resolution-aware mapping between experimental MS features and database entities. Importantly, this architecture allows OmniPath Metabo to accommodate the intrinsic uncertainty of metabolomics annotations rather than forcing all entities into a single flat identifier namespace.

Our results further illustrate that metabolomics interpretation benefits from moving beyond conventional pathway enrichment approaches. In the lung cancer case study, differential metabolites such as azelaic acid could not be adequately interpreted through pathway membership alone. Instead, biologically relevant context emerged from combining chemical annotations, disease associations, literature links, and metabolite-protein interaction knowledge. This highlights an important conceptual point: metabolites are not merely intermediates within metabolic pathways, but active participants in signaling, transport, redox regulation, and intercellular communication. OmniPath Metabo was specifically designed to capture this broader functional landscape. This metabolite-centered perspective also motivates our ongoing efforts toward a more biologically realistic representation of cellular regulation.

By integrating receptor-mediated signaling, transporter activity, and allosteric regulation, tools such as COSMOS+ [33] or CORNETO [44] become able to model metabolites as active causal entities that can influence intracellular signaling and transcriptional programs. This is especially relevant for biological contexts where extracellular metabolites mediate communication between cells or tissues, including cancer metabolism, immunometabolism, and microbiome-host interactions.

Although the current release already integrates 40+ resources, OmniPath Metabo is designed as an extensible platform rather than a static database. A major future direction will be the incorporation of additional resource categories that connect metabolism to broader biological ecosystems. One particularly important area is microbiome-associated metabolism. Microbial metabolites are increasingly recognized as key mediators of host physiology, immunity, and disease, yet microbiome-metabolite-host interaction knowledge remains highly fragmented across specialized resources. While this functionality is not yet included in the current preprint release, the modular architecture of the database was explicitly designed to support such expansions.

In summary, OmniPath Metabo transforms fragmented metabolomics and systems biology resources into a unified, structure- and ontology-aware, and mechanism oriented knowledge framework. The framework not only supports exploratory data interpretation through its web interface and API, but also serves as a generator of application-specific prior knowledge subsets. Subsets such as the prior knowledge network of multi-omics metabolite-driven regulation (COSMOS [34, 45] related network) aim to provide the foundation for next-generation multi-omics and spatial omics modeling of multi-omics metabolite-driven regulation. Similar specialized subsets coming up soon will cover 1) cell-cell communication; 2) drug-target interactions; 3) lipid signaling and metabolism. We anticipate that OmniPath Metabo will facilitate more reproducible, interoperable, and mechanistically informative metabolomics analyses across a wide range of biomedical applications. As this is an ongoing effort, we want to note that the current database and results will be subject to frequent updates. We do our best to assure thorough quality control and stabilization of the design, yet users should take into account that this is an early phase of the project. We share it with the community in this early phase to collect feedback.

## 4. Methods

### 4.1. OmniPath Metabo Resources

### 4.2. Build pipeline and data integration

The OmniPath Metabo database is built through a modular pipeline that separatesdataacquisition, resource-specificnormalization, cross-resourceintegration through entity resolution, and final materialization for the serving layer.

#### 4.2.1. Input Modules

Source data enter the pipeline through ‘input‘ modules. Each module defines one or more datasets and provides the source-specific knowledge needed to retrieve, parse, and interpret those datasets. At the ingestion level, each external resource is represented declaratively as one or more datasets. A dataset combines three components: (i) a download specification describing how the source is retrieved, (ii) a raw parser that converts source files into record dictionaries, and (iii) a schema mapper that transforms those records into the shared internal ‘Entity‘ representation.

This design allows heterogeneous upstream formats to be processed through a common ingestion model. Input modules are therefore not only responsible for parsing source files, but also for defining how source-specific fields can be mapped to the corresponding set of identifier namespaces, annotation terms, entity types, and other controlled vocabularies.

#### 4.2.2. Entity Representation

Resource-level normalization is performed against an internal ontology-backed schema. Each normalized ‘Entity‘ consists of four core elements: (i) an entity type, (ii) a set of typed identifiers, (iii) a set of typed annotations, and (iv) optional membership relations to other entities. This allows proteins, metabolites, interactions, complexes, pathways, reactions, and ontology terms to be represented with the same structural model.

Entity fields are encoded using ontology terms, primarily from the Proteomics Standards Initiative Molecular Interactions ontology [46], supplemented where needed with OmniPath-specific terms. As a result, identifiers, and annotations are normalized to explicit semantic terms during ingestion.

#### 4.2.3. Source Evidence Projection

For each selected dataset, the pipeline executes the corresponding raw parser and streams the emitted records into the build process. Parsed records are not loaded directly as final graph records. Instead, they are first projected into source-scoped evidence tables in DuckDB [47]. Each evidence row records: the source, dataset, source row position, entity occurrence path, identifiers, annotations, and membership or interaction structure.

Evidence identifiers are generated deterministically from the source, dataset, row identifier, and occurrence path. This provides stable evidence keys for repeated builds over the same parser output and preserves the boundary between source-specific observations and canonical graph entities. Entity records, interaction-like records, membership structures, and ontology-derived annotations are all flattened into a shared evidence representation before canonicalization.

#### 4.2.4. Entity canonicalization

Protein and gene entities are preferentially resolved to their corresponding Entrez/NCBI Gene identifiers, while chemical entities are preferentially resolved to standardized InChI Keys. The chemical resolver lookup materializes authoritative issuer mappings, such as those provided by ChEBI [18], HMDB [26], LipidMaps [29], SwissLipids [23], Pubchem [28], and RefMet [31].

For each evidence entity, the pipeline collects all identifiers and queries the resolver lookup tables for accepted canonical candidates. Chemicals which cannot be mapped via the resolver lookup to a structure are resolved through a fallback hierarchy that deterministically selects a supported identifier available for that evidence record. Direct canonical identifiers are preferred over authoritative issuer mappings, which are preferred over the fallback hierarchy.

A single top-ranked candidate resolves the evidence entity to the corresponding canonical entity. If fallback also fails, the entity is retained as unresolved with a stable fallback identifier derived from its identifier set, entity type, and taxonomy context where applicable. Multiple equally ranked candidates are marked as ambiguous and are handled like the unresolved case. The resolution workflow is summarized in Figure 6. Detailed per-resource identifier coverage is provided in Supplementary Table 2.

**Figure 6.**
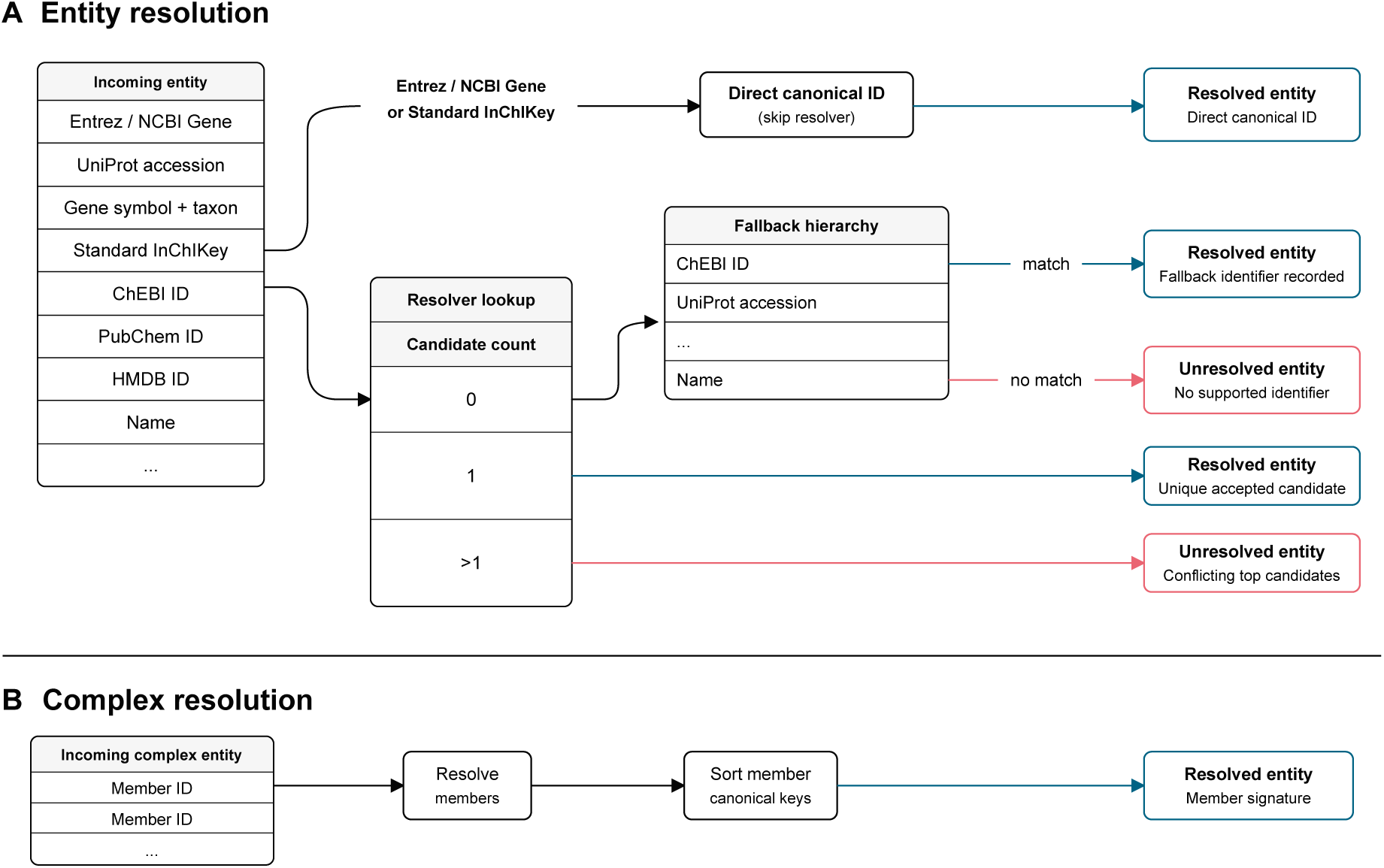
Entity and complex resolution workflow. Incoming entities with Entrez / NCBI Gene identifiers or standard InChIKeys already carry preferred canonical identifiers and therefore bypass resolver lookup. Other identifiers are queried against evidence-independentresolver lookup tables. A single accepted candidate resolves the entity; no accepted candidate triggers the fallback hierarchy; and multiple equally supported candidates mark the entity as ambiguous and leave it unresolved. Complexes are resolved by resolving their members and deriving a sorted canonical member signature.

#### 4.2.5. Provenance and Reproducibility

The final database contains two connected data layers: source-scoped evidence anddeduplicatedgraphrecords. Evidencetablespreservethenormalizedinter-pretation of each parsed source row, including source, dataset, row identifier, entity occurrence path, identifiers, annotations, and relation evidence. Graph tables provide canonical entities and relations for query and integration. Every graph entity or relation can be traced back through evidence links to its source, dataset, and source row. This preserves source-level provenance while allowing equivalent records from different datasets to collapse into shared graph entities and relations.

The pipeline is incremental at the source level. Existing source content can be skipped, refreshed, or deleted independently. On reload, stale evidence for the selected source is removed before current parser output is projected and canonicalized again. Canonical graph rows that remain supported by other sources are preserved, while unsupported source-specific evidence is removed. This allows selected sources to be refreshed without rebuilding the full database.

### 4.3. Prior knowledge network of multi-omics metabolite-driven regulation

#### 4.3.1. Network architecture

To modelmechanistichypothesesspanningsignalingpathwayandmetabolism, we constructed a prior knowledge network (PKN) by connecting six types of directed, signed interactions into a unified network: transporter-metabolite, receptor-metabolite, allosteric regulation (metabolite-enzyme protein), metabolic reaction (enzyme-metabolite), protein-protein interaction (PPI), and transcription factor-target gene regulation Gene Regulatory Network (GRN) (Table 3).

#### 4.3.2. ID translation

To unify identifiers across all resources, metabolite IDs were translated to ChEBI ID [18, 19] and protein IDs to UniProt accessions [48] using resource-specific strategies. From PubChem [28] to ChEBI [18, 19], we used the combined UniChem [49]/RaMP [4]/MetaNetX [50] mapping tables (182,000 pairs). From MetAtlas IDs [50] to ChEBI [18, 19], we applied a six-step fallback strategy: (i) direct annotation in GEM metabolite table, (ii) MetaNetX [50], (iii) LipidMaps via UniChem [49], (iv) PubChem via UniChem [49], (v) KEGG REST [5], (vi) HMDB [26, 27]. From metabolite name (synonym) to ChEBI, we applied a three-step name-lookup strategy: (i) HMDB [26, 27], (ii) RaMP [4], (iii) PubChem REST [28]. From the Ensembl gene or protein to Uniprot, we translated via omnipath-utils mapping API. The remaining resources were directly pass-through as they are natively using UniProt or ChEBI.

#### 4.3.3. Metabolite–protein interactions: transporters, receptors, and allosteric regulations

We assembled transporter-metabolite interactions from five resources (Table 3). For TCDB [51, 52], we filtered interactions to the target organism by matching transporter UniProt accessions against the organism-specific UniProt reference list. For MRCLinksdb [53], we annotated each protein as transporter or receptor by using Guide to Pharmacology [54, 55] [54], TCDB [51, 52], and OmniPath Intercell [15, 56]. From the Metabolic Atlas [41, 57], we identified transport-metabolite interactions by the subsystem annotation ‘Transport reactions’; reactions without a gene rule were retained as orphan pseudo-enzyme nodes. From the BiGG genome-scale metabolic models [40], we identified transport reactions by compartment-crossing detection, retaining any reaction in which the same metabolite identifier appeared in two different compartments.

We assembled receptor-metabolite interactions from MRCLinksdb [53] and STITCH [58]. For STITCH [58], we annotated each protein using the same schema as MRCLinksdb described in the last paragraph. The rest procedures followed our previous work [34].

We obtained allosteric regulations from BRENDA [42] following the methodology that has been reported previously [59, 60].

#### 4.3.4. Metabolite–enzyme interactions

We obtained metabolite-enzyme interactions from Metabolic Atlas genome-scale models [41, 57] and BiGG models [40], supplemented with enzyme-metabolite edges from KEGG retrieved via the KEGG REST API [5]. Each stoichiometric reaction was decomposed into directed substrate and enzyme, enzyme and product binary edges; reversible reactions additionally produce the reverse-direction pair.

Cofactor handling differs by source. For BiGG models [40], a degree-based filter removes metabolites appearing in more than 400 edges across the network, excluding ubiquitous cofactors such as ATP and NADH that would otherwise generate spurious high-degree hubs. Metabolic Atlas [41, 57] GEM and KEGG [5] apply no cofactor filter. Reactions with no gene annotation are handled differently per source: Metabolic Atlas [41, 57] retains orphan reactions as pseudo-enzyme nodes; BiGG models [40] and KEGG [5] skip enzyme-free reactions.

#### 4.3.5. PPI and GRN

PPI and GRN were retrieved from the OmniPath web service, retaining only signed, directedinteractions(is_stimulationoris_inhibitionequalto 1). Where both stimulatory and inhibitory evidence were reported for the same node pair, we preserved each as a separate edge.

#### 4.3.6. Cellular location

For TCDB [51, 52] and MRCLinksdb [53], we annotate subcellular location information by UniProt [48]. For SLC TABLES [61], compartment information comes directly from the SLC TABLES database’s own localization field [61]. For GEM models (Metabolic Atlas [41, 57] and BiGG models [40]), compartment codes are embedded in the model stoichiometry as single-letter metabolite suffixes and are used directly without further annotation.

All compartment labels are normalised to the following abbreviations, aligned with BiGG model conventions [40]. e: extracellular/plasma membrane; n: nucleus; c: cytoplasm; m: mitochondrion; r: endoplasmic reticulum; g: golgi apparatus; eg: endoplasmic reticulum-Golgi; v: vesicle; l: lysosome; x: peroxisome. Resources without explicit compartment information (KEGG [5], STITCH [58], BRENDA [62], and OmniPath signaling resources) carry no location annotation in the PKN.

#### 4.3.7. Network assembly, formatting, and availability

Transporter: For TCDB [51, 52], SLC TABLES [61], and MRCLinksdb [53], which provide a single substrate and transporter edge, we expanded each interaction into four directed edges during formatting. For Metabolic Atlas [41, 57], BiGG model [40], and KEGG [5], which already provide directional reaction edges.

The 6 interaction layers were combined into a single network. Within each category, duplicate edges sharing the same source, target, and MOR were merged by unioning compartment annotations. Network nodes were reformatted as follows. Network nodes were reformatted as follows: Metab CHEBI:ID_<compartment>; transporter and GEM enzyme nodes as Gene{N} UniProtAC (forward), and Gene{N} UniProtAC_rev (reverse), where N is a sequential integer per unique reaction within each category; orphan pseudo-enzyme nodes as Gene{N} orphanReac<reaction_id=. Receptors, allosteric targets, and all signaling layer proteins (PPI and GRN) use bare UniProt accessions. Connector edges (bare UniProtAC to Gene{N} UniProtAC) were appended to bridge the signaling layer to the metabolic layer, but only for proteins that also appear as nodes in the PPI, GRN, or ligand–receptor sub-networks.

The complete COSMOS+ PKN build pipeline is implemented in the open-source omnipathmetabo python package (https://github.com/saezlab/omnipath-metabo) and is programmatically accessible via a REST API at metabo.omnipathdb.org.

### 4.4. Cancer cell line metabolomics data

#### 4.4.1. Preprocessing of metabolomics data

Metabolomics data from Shorthouse et al. 2022 [35] was imported from the supplementary excel files containing sample-level metadata, metabolite feature annotations, and ion intensity measurements. For samples, the reported cell line name and associated descriptive metadata fields were retained, while repeated values within each injection were collapsed into delimited character strings to preserve all source annotations. Placeholder values such as ‘NA‘ and ‘NS‘ were standardized to missing values.

Because the downstream biological analyses were designed at the cell line level, technical replicate injections were collapsed by cell line identity. In parallel, the ion annotation table was cleaned by removing summary columns not required for downstream analysis and by replacing spaces in metabolite names with underscores for stable string handling. An additional combined identifier field (‘id_list‘) was constructed from non-missing HMDB and KEGG identifiers to support duplicate detection.

Duplicate metabolite features were identified using the combined identifier field and removed before quantitative processing. This step specifically addressed cases where different ion m/z entries had been assigned the same metabolite name and HMDB identifier, for example through alternative adduct representation. Intensity values were then extracted from the quantitative matrix, collapsed across technical replicates and the arithmetic mean was calculated for each metabolite feature across replicate injections. The resulting collapsed dataset consisted of a cell line-level intensity matrix together with matched sample and feature metadata. A detailed script can be found at https://github.com/saezlab/metabo-usecases/tree/main/omnipath_metabo_case1/scripts.

#### 4.4.2. Cell line metadata enrichment

The collapsed Shorthouse cell line object was enriched with external biological metadata derived from Cellosaurus [63]. The Cellosaurus flat file [https://ftp.expasy.org/databases/cellosaurus/cellosaurus.txt, accessed on 19.05.2026] was parsed into an entry-level lookup table in which each accession was linked to its recommended cell line name and all available synonyms. Matching between Shorthouse cell line names and Cellosaurus entries was performed by joining the reported cell line name against this synonym-expanded lookup. Names that could not be resolved automatically were curated manually, and ambiguous mappings were resolved using a manually defined preferred-accession table for selected cell lines with multiple accession matches.

Once a preferred Cellosaurus accession had been assigned to each Shorthouse cell line, the Cellosaurus API was queried to retrieve accession-linked meta-data fields including sequence variation, cell type, and derived anatomical site. Mutation content was extracted from the Cellosaurus ‘sequence-variation‘ field. These mutation annotations were then joined back onto the Shorthouse sample metadata together with the derived anatomical site, and Cellosaurus cell type. The resulting accession mapping was cross-checked against the Research Resource Identifier (RRID) Cellosaurus accessions which were already present in the Shorthouse metadata, concluding a complete match and increased coverage through our approach.

Metabolite feature metadata were additionally enriched with broad chemical class information using the MetaProViz ‘metsigdb_chemicalclass()‘ prior knowledge retrieval [64]. To support tissue-level stratification of cancer cell lines, sample origin was harmonized into a generalized anatomical site variable. Two complementary rule-based mapping schemes were implemented. One was based on disease descriptors and one based on the Cellosaurus-derived anatomical source site. The sample site information in Cellosaurus can reflect metastatic fluid or culture source material rather than the underlying tumour lineage. Because we were interested in disease classification rather than sampling sites, disease-driven mapping was prioritized. The final ‘general_site‘ variable was defined by coalescing the disease-based and site-based assignments. The final enriched Shorthouse data comprised metabolite intensities, feature annotations, Cellosaurus-derived mutation metadata, chemical class assignments, and harmonized tissue labels. A detailed script can be found at https://github.com/saezlab/metabo-usecases/tree/main/omnipath_metabo_case1/scripts.

#### 4.4.3. Metabolite identifier quality control, expansion, and reconstruction of the extended feature set

Metabolite feature processing was performed exclusively on the feature metadata table of the annotated Shorthouse data and was designed to improve the completeness and internal consistency of small-molecule identifiers across HMDB, KEGG, ChEBI, and PubChem. Identifier completeness and multiplicity of the original feature table were quantified independently for each database using MetaProViz ‘count_id()‘ [64] , which classifies features as having no identifier, a single identifier, or multiple identifiers. MetaProViz ‘compare_pk()‘ [64] was used to summarize overlap across identifier systems. These summaries established the baseline metabolite annotation state before curation. Internal consistency of seed identifiers (=IDs provided by original authors) was then assessed using the MetaProViz seed identifier compatibility workflow [64]. Pairwise compatibility among HMDB, KEGG, ChEBI, and PubChem assignments was evaluated for each feature, and features were classified as fully compatible, partially compatible, or completely incompatible. Features classified as partially compatible were filtered to retain only identifier relationships supported by direct or secondary matching paths in the ID graph, whereas incompatible relationships marked as ‘no_match‘ were discarded. Fully compatible feature annotations were retained unchanged. Metabolite identifiers were expanded across HMDB, KEGG, ChEBI and Pub-Chem namespaces by a similar graph-based traversal across identifier systems using MetaProViz ‘traverse_ids()‘ [64]. To avoid uncontrolled propagation through excessively large mapping connections in the ID graph, translated identifiers were only accepted when the number of translated identifiers for a given database remained below 50. The traversed feature table was again summarized with ‘count_id()‘ and ‘compare_pk()‘ to assess changes introduced by network traversal relative to both the original and cleaned states. As a final identifier enrichment step, MetaProViz ‘equivalent_id()‘ [64] was applied independently to HMDB, KEGG, ChEBI, and PubChem identifiers to add database-equivalent metabolite identifiers not already captured by the traversal step. This added stereochemically equivalent IDs across namespaces for downstream analyses, representing the final version of the feature metadata. A detailed script can be found at https://github.com/saezlab/metabo-usecases/tree/main/omnipath_metabo_case1/scripts.

#### 4.4.4. Differential metabolite analysis and mutation pattern assessment

Differential metabolite abundance analysis was performed for Shorthouse [35] data, restricted to cell lines annotated as originating from “Lung/Pleura”, as these represented ∼ 45 % or cell lines and hence the largest population in the dataset. Because oncogenic drivers in these cell lines frequently co-occur with additional mutations, we analysed co-occurence of mutations for the following defined contrasts.

Four pairwise comparisons were defined: (i) KRAS-mutant versus KRAS-wild-type lung cancer cell, (ii) KRAS-mutant versus KRAS-wild-type lung cancer cell further restricted to TP53-mutant, EGFR-wild-type, CDKN2A-wild-type, and STK11-wild-type backgrounds, (iii) EGFR-mutant versus EGFR-wild- type lung cancer cell lines, and (iv) EGFR-mutant versus EGFR-wild-type lung cancer cell lines further restricted to TP53-mutant, KRAS-wild-type, CDKN2A-wild-type, and STK11-wild-type backgrounds. This design was intended to capture both the overall metabolic signal associated with KRAS or EGFR status and a more specific driver-associated signal after reducing potential confounding by major co-mutation patterns.

Feature metadata were aligned to the intensity matrix for filtering and grouping based on these pre-defined contrasts. Where available, metabolite names and metabolite-class labels were propagated from the feature annotation table for labelling and visualization. Intensity values were filtered before logarithmic transformation to avoid propagating negative values into the linear model.

Differential analysis was carried out with the “limma” framework [65]. Per comparison, all samples were assigned to a two-level factor representing mutation-positive and mutation-negative groups, with the mutation-negative group used as the reference level. For each retained metabolite, “limma::lmFit”, “limma::contrasts.fit”, and “limma::eBayes” were applied sequentially to estimate moderated t-statistics, raw p-values, and empirical Bayes-shrunken standard errors. P-values were adjusted for multiple testing within each comparison using the Benjamini-Hochberg false discovery rate procedure [66].

To support interpretation of the mutational background represented in each cohort, the script additionally reported the frequency of all mutation tokens among all samples in a comparison and among mutation-positive samples only. Outputs of the differential analysis and mutation profiles of the samples used in this study are provided in **Supplementary Table 4**, which was generated using the corresponding scripts available in the repository (https://github.com/saezlab/metabo-usecases/tree/main/omnipath_metabo_case1/scripts).

#### 4.4.5. Extracting Azelaic acid information

Azelaic acid annotations and interactions were extracted from the OmniPath Metabo using the FastAPI endpoints (https://dev.omnipathdb.org/api/docs). Candidate entities were obtained by querying with with GET /entities/search for “Azelaic acid (PTF52203)”, “Azelaic acid; nonanedioic acid”, and “Azelate”, and cross-checked with POST /entities/resolve, providing the retrieved identifiers. The first two entities are unresolved records, respectively from PTFI and Recon3D, while the latter is the resolved Azelate entity, across multiple original data sources, identified by Standard InChIKey BDJRBEYXGGNYIS-UHFFFAOYSA-N.

Selected entities were hydrated with POST /entities/by-pks to retrieve their annotations; direct relation records were retrieved for each seed with paginated GET /relations/search; and for each retrieved relationPk, evidence was fetched separately via GET /relations/{relationPk}/evidence. All the information retrieved by FastAPI queries, is available in **Supplementary Tables 5**, including seed entity resolution and identifiers, source memberships, direct relations and evidence counts, HMDB/ChemOnt class links, FooDB food-occurrence links, metabolic reactions and transport annotations, and MACdb disease-association evidence.

For plotting, only records linked to the resolved “Azelate” entity were retained. Interaction summaries were generated by separating relations annotated with multiple sources into one row per source, then counting each unique relationPk once within each interaction type and source.

Cancer-association summaries were generated from MACdb evidence only. Records were first filtered to disease terms containing cancer-related keywords, then sample labels were harmonized by grouping tissue-specific labels as “Tissue” and renaming “A2780 cells” as “Cells”. To avoid counting the same study multiple times, duplicate rows with the same disease type, sample type, PubMed/PMC identifier, study description, case description, and control description were collapsed before counting evidence records per disease type and sample type.

**Supplementary Table 5** outputs were created using a collection of scripts in https://github.com/saezlab/metabo-usecases/tree/main/omnipath_metabo_case1/azelate_fig_tables/notebooks.

#### 4.4.6. Linking altered metabolites to GEMs, allosteric regulations and cellular location

For each EGFR- and KRAS-mutant comparison, we selected the 10 metabolites with the highest and the 10 with the lowest moderated t-statistics from the differential analysis, resulting in 20 metabolites per comparison without applying an additional significance threshold. Metabolites were classified as up- or down-regulated according to the sign of their log-fold change. Where multiple ChEBIidentifierswereassignedtoametabolite, theseidentifierswere separated and considered individually.

The selected ChEBI identifiers were mapped to the COSMOS PKN [34, 45]. We considered two interaction layers: enzyme–metabolite relationships supported by KEGG [5], Human-GEM [57], and Recon3D [67], with orphan metabolic reactions excluded; and metabolite–protein regulatory relationships supported by BRENDA [42] and STITCH [58]. Evidence from multiple resources was retained when available. An interaction was selected when either its source or target matched one of the altered metabolites, thereby capturing all directly connected PKN edges rather than only downstream interactions.

Matched interactions were summarized separately for EGFR and KRAS, metabolite direction, interaction layer, and contributing resource. Subcellular locations were obtained from the compartment annotations associated with the PKN edges and included the cytoplasm, mitochondria, nucleus, extracellular space, endoplasmic reticulum, peroxisome, Golgi apparatus, vesicles, and lysosomes. Interactions assigned to multiple compartments were counted in each annotated compartment, whereas interactions without compartment information were omitted from the location-specific analysis.

## Code and Data availability

An integrated pipeline for generating figures, tables and case studies of this paper can be found in the https://github.com/saezlab/metabo-usecases repository. Data of the main OmniPath database is available at https://dev.omnipathdb.org, the Utils database at https://utils.omnipathdb.org/ and the Metabo API at https://metabo.omnipathdb.org/. The major soft-ware packages presented in this paper are available at https://github.com/saezlab/pypath (omnipath-resources; resource access and pars-ing); https://github.com/saezlab/omnipath-build (main database build); https://github.com/saezlab/omnipath-present (web application); https://github.com/saezlab/omnipath-utils (OmniPath Utils database build and service); https://github.com/saezlab/omnipath-metabo (OmniPath Metabo build extension and web API); https://github.com/saezlab/omnipath-client (Python client for all OmniPath APIs).

## Acknowledgements

We thank all members of the Saez Lab, specifically Forrest Hyde and Akane Ogawa, for valuable discussions and feedback throughout the development of OmniPath Metabo. We are particularly grateful to the members of the Omni-Path development team for their contributions to database development, software infrastructure, and resource integration. We thank Dr. Keigo Morita from the Graduate School of Science, University of Tokyo, for his assistance with the implementation and use of the BRENDA API.

We also acknowledge the developers and maintainers of the numerous public databases and resources integrated into OmniPath Metabo. Their continuous efforts in generating, curating, and maintaining biological knowledge make integrative resources such as OmniPath Metabo possible.

J.S. and J.SR. acknowledge funding from the European Union’s Horizon 2020 Programme under grant agreement no. 965193 (DECIDER, https://www.deciderproject.eu/); Y.B. from the Medical Data Scientist Program of the Heidelberg Faculty of Medicine; T.L. from the UKRI BBSRC [BB/Y512540/1]; N.PE. from the Deutsche Forschungsgemeinschaft (DFG, German Research Foundation) through SFB 1550 (DFG-508152189); M.D. from SmartCare (03LW0233K); T.K. from the NIHR Imperial Biomedical Research Centre Organoid Facility; B.B., L.G. and T.K. from the UKRI BBSRC Institute Strategic Programme Food Microbiome and Health BB/X011054/1 and its constituent project BBS/E/F/000PR13631; E.C. from the Deutsche Forschungsgemeinschaft (DFG) grant number 528753569; D.M. from the European Union’s Recovery and Resilience Facility-Next Generation, in the framework of the General Invitation of the Spanish Government’s public business entity Red.es to participate in the talent attraction and retention programmes within Investment 4 of Component 19 of the Recovery, Transformation and Resilience Plan (SOLI/2024/0524/00240212); C.S. from the LiSyM-cancer network supported by the German Federal Ministry of Education and Research (BMBF) (031L0257B); D.T. from the Landesinstitut für Bioinformatikinfrastruktur in Baden-Württemberg.

## Contribution Notes

J.S.: Writing original draft; software/technical implementation; OmniPath Metabo schema and backend; website development.

Y.B.: Writing original draft; figure creation; development of Cosmos+ PKN.

J.F.: Writing original draft; figure creation; software/technical implementation; use-case data acquisition and analysis.

T.L.: Writing original draft; figure creation; software/technical implementation; development of protein-metabolite interactions.

N.PE.: Software/technical implementation

D.B. Writing original draft; figure creation; software/technical implementation; database searches for use cases; website development.

E.C.: Ontology development.

M.D.: Preliminary software/technical implementation of PKN generation from lipid databases (SwissLipids, Rhea, Reactome, LipidMaps).

L.G.: Supervision; conceptualization; software/technical implementation

A.S.: Figure creation; software/technical implementation - KEGG.

D.M.: Software/technical implementation - preliminary Recon3D and GEMS scripts forming the basis for Cosmos+ PKN.

B.B.: Software/technical implementation

A.D.: Software/technical implementation - preliminary Recon3D and GEMS scripts forming the basis for Cosmos+ PKN.

T.K.: Supervision; conceptualization.

D.T.: Figure creation; supervision; conceptualization; project development; software/technical implementation; development of Cosmos+ PKN; Omni-Path Metabo schema and backend; website development.

C.S.: Writing original draft; figure creation; supervision; conceptualization; project development; use-case and metabolism-related work.

J.SR.: Supervision; conceptualization. Everyone: Revising manuscript.

## Conflict of interests

JSR reports in the last 3 years funding from GSK and Pfizer & fees/honoraria from Travere Therapeutics, Stadapharm, Astex Pharmaceuticals, Owkin, Pfizer, Vera Therapeutics, Grunenthal, Tempus and Moderna

## Supplementary Figures

**Supplementary Figure S1.**
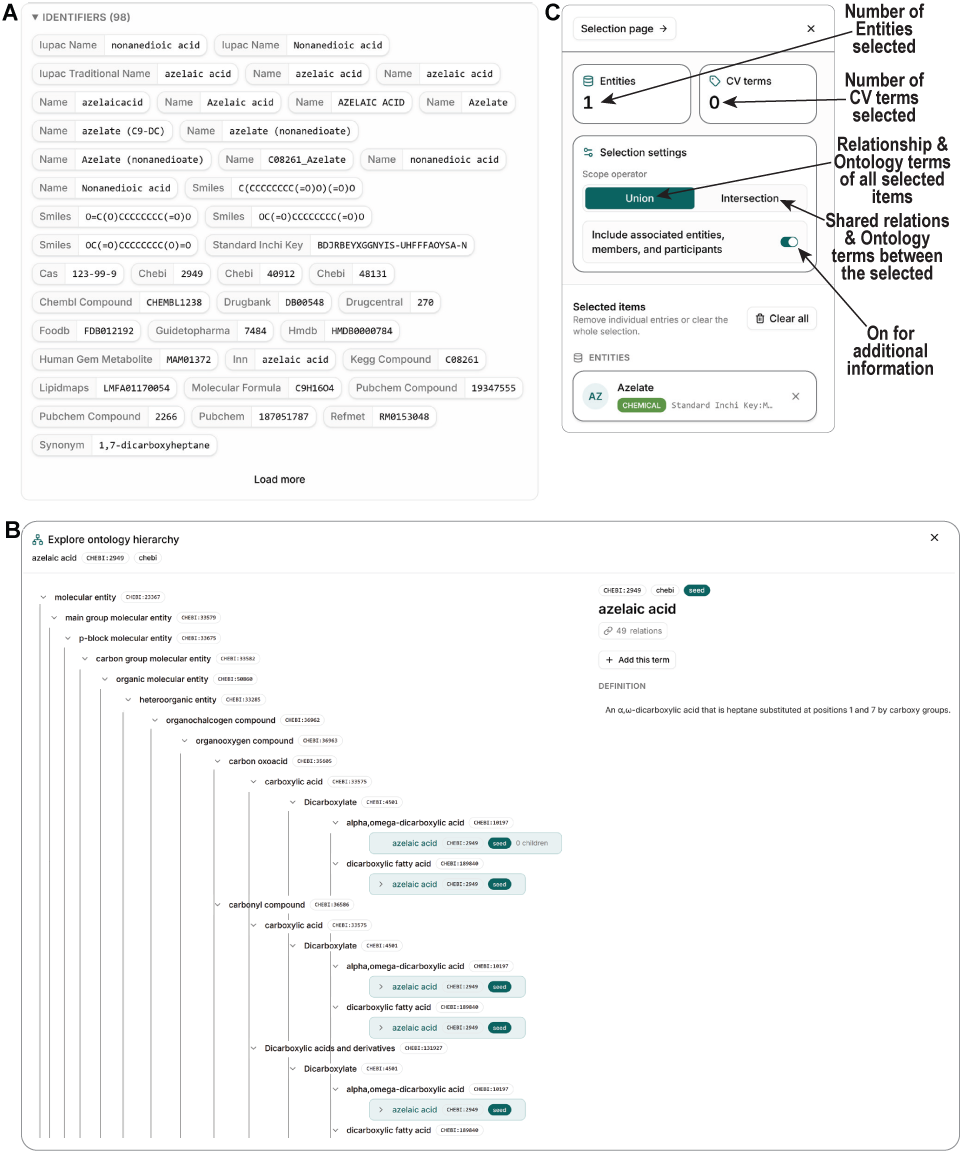
Additional functionalities of the OmniPath Metabo web interface. **A**) Expanded identifiers view from the entity tab showing detailed feature identifiers such as trivial names, chemical structures (i.e. Smiles) and database identifiers (i.e. ChEBI IDs) amongst other feature identifiers. **B**) Hierarchy and ontology browser enabling navigation of parent-child relationships, ontology terms, and associated biological entities. Here the ontology hierarchy for azelaic acid is displayed. **C**) Selection workspace illustrating context-specific exploration of user-selected entities and annotations. Selected entities can be used to retrieve associated relations and ontology terms, while ontology terms can be used to iteratively refine entity selections and identify biologically coherent subsets.

**Supplementary Figure S2.**
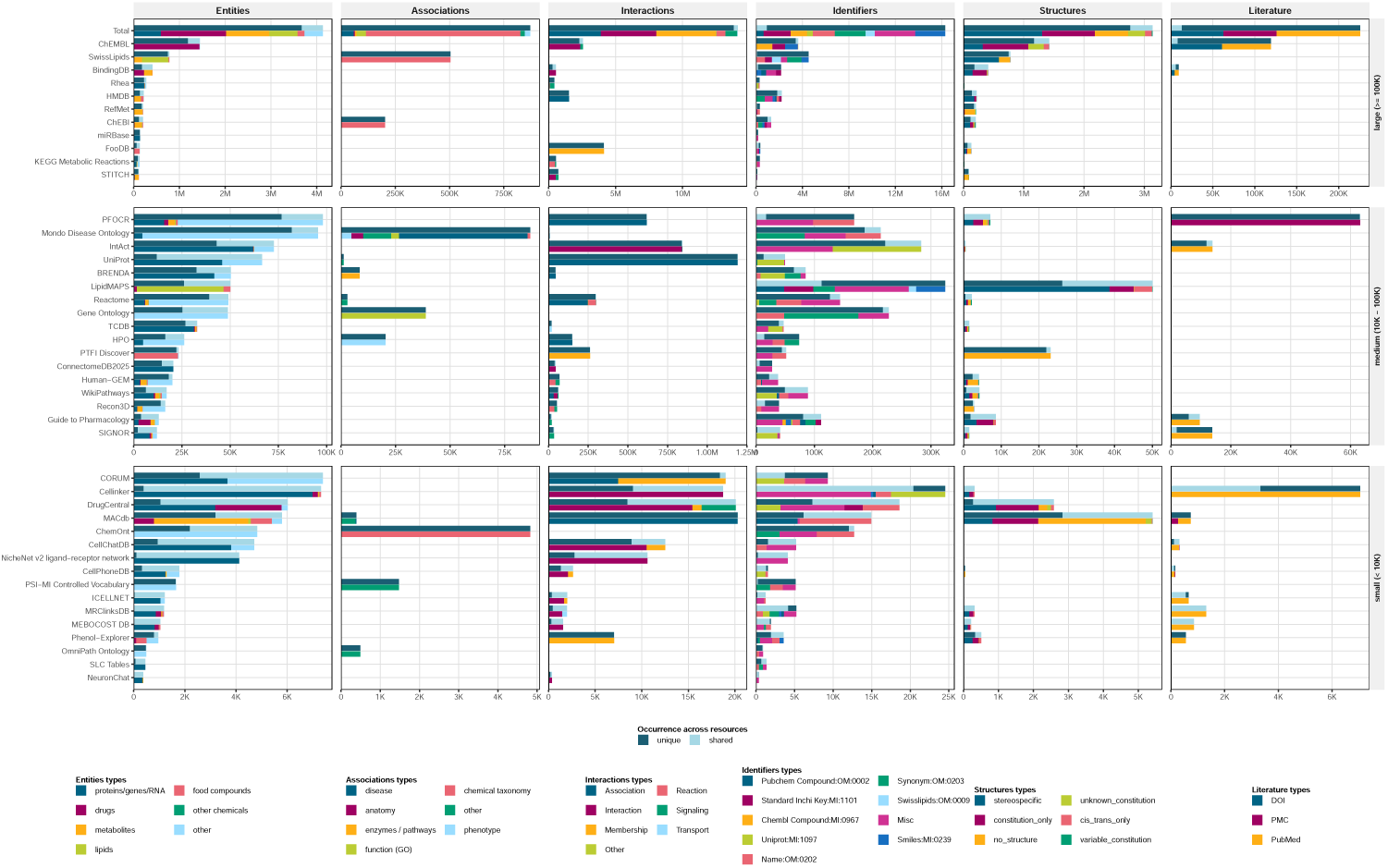
OmniPath Metabo Database Content. Horizontal bar charts summarize the content contributed by each source database, grouped by resource size (large, >100K records; medium, 10K–100K records; small, <10K records). Columns represent the major data categories integrated into the database: Entities, Associations, Interactions, Identifiers, Structures, and Literature. Bar lengths indicate the number of records contributed by each resource. Colored segments denote data subtypes within each category (e.g., proteins/genes/RNAs, drugs, metabolites, lipids, diseases, phenotypes, signaling interactions, chemical identifiers, molecular structures, and literature references), as indicated in the legends. For entities, interactions, identifiers, and structures, bars are further partitioned into records that are unique to a given resource versus those shared across multiple resources.

## Supplementary Tables

**Supplementary Table 1.**
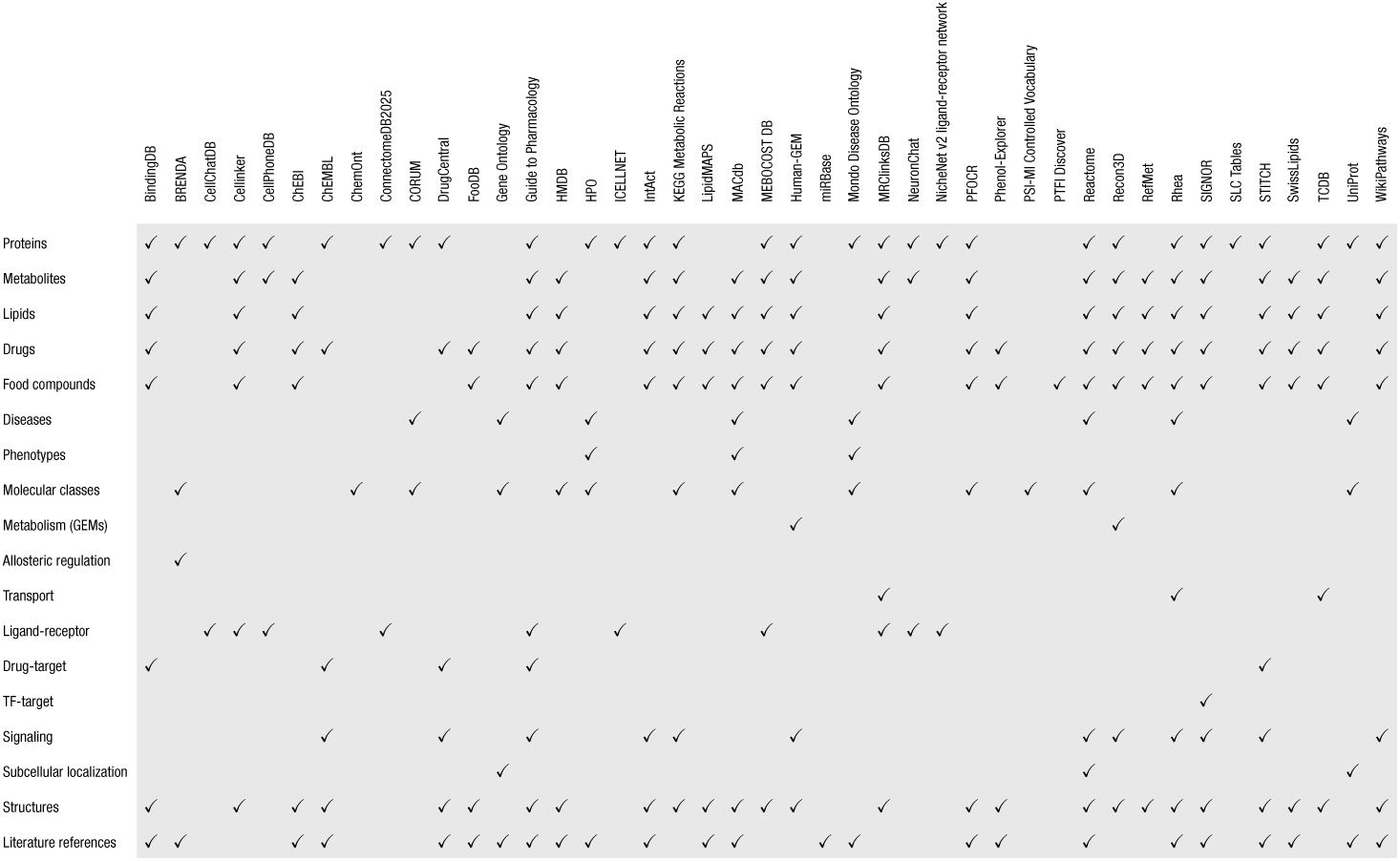
Content coverage of integrated source databases. Overview of biological entities, molecular classes, interaction types, structuralinformation, andliteratureannotationscontributedbyeachsourcedatabase. Checkmarksindicatewhetheraresourceprovidesdata for a given category.

**Supplementary Table 2.**
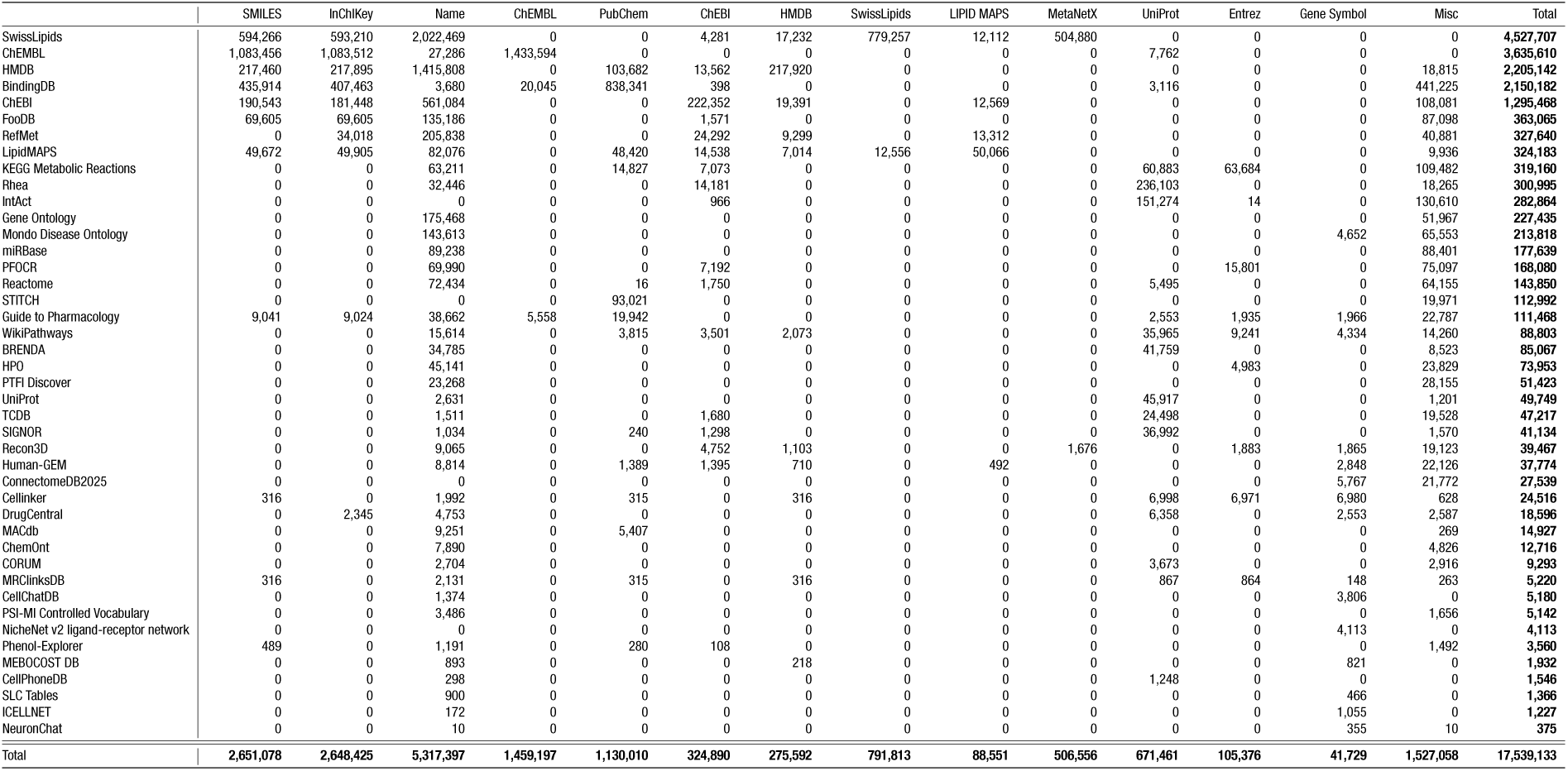
Identifier coverage across integrated source databases. Coverage of identifier types provided by each integrated resource and incorporated into the database. Rows correspond to source databases and columns represent identifier categories, including chemical structure descriptors (SMILES and InChIKeys), names, database-specific identifiers (e.g., ChEMBL, PubChem, ChEBI, HMDB, SwissLipids, LIPID MAPS, MetaNetX, UniProt, and Entrez Gene), gene symbols, and miscellaneous cross-references. Values indicate the number of records contributed by each source for each identifier category, with totals summarizing the overall identifier content integrated from each resource.

**Supplementary Table 3.** Characterization of ambiguous compound mappings from RampDB. Conflict categories include mappings differing only by stereochemistry, mappings representing different levels of chemical specificity, unrelated structures, closely related structures, and tautomeric variants. For each category, the table reports the number of distinct database identifiers involved, the number of conflicting identifier pairs, the percentage contribution to the total set of conflicts, and concrete examples.

|  | Distinct RaMP ids | Conflict pairs | Share (%) | Example RaMP ids |
| --- | --- | --- | --- | --- |
| Stereochemistry only | 20,731 | 21,847 | 92.5 | RAMP_C_000000002, RAMP_C_000000003 |
| Different specificity levels | 1,315 | 1,778 | 5.9 | RAMP_C_000000004, RAMP_C_000000027 |
| Unrelated structures | 267 | 452 | 1.2 | RAMP_C_000000001, RAMP_C_000000045 |
| Similar structures | 74 | 132 | 0.3 | RAMP_C_000000410, RAMP_C_000000446 |
| Tautomeric variants | 29 | 43 | 0.1 | RAMP_C_000000517, RAMP_C_000000646 |
| **Total** | **22,416** | **24,252** |  |  |

## Notes

https://github.com/saezlab/metabo-usecases

https://utils.omnipathdb.org/

https://metabo.omnipathdb.org/

https://github.com/saezlab/pypath

https://github.com/saezlab/omnipath-build

https://github.com/saezlab/omnipath-present

https://github.com/saezlab/omnipath-utils

https://github.com/saezlab/omnipath-metabo

https://github.com/saezlab/omnipath-client

## References

[1] Consistency, Inconsistency, and Ambiguity of Metabolite Names in Biochemical Databases Used for Genome-Scale Metabolic Modelling. https://www.mdpi.com/2218-1989/9/2/28.

[2] Romanas Chaleckis, Isabel Meister, Pei Zhang, and Craig E Wheelock. Challenges, progress and promises of metabolite annotation for LC–MS-based metabolomics. Current Opinion in Biotechnology, 55:44–50, February 2019. ISSN 0958-1669. doi: 10.1016/j.copbio.2018.07.010.

[3] Carlos Calderón and Michael Lämmerhofer. Enantioselective metabolomics by liquid chromatography-mass spectrometry. Journal of Pharmaceutical and Biomedical Analysis, 207:114430, January 2022. ISSN 0731-7085. doi: 10.1016/j.jpba.2021.114430.

[4] J Braisted, A Patt, C Tindall, T Sheils, J Neyra, K Spencer, T Eicher, and EA Mathé. RaMP-DB 2.0: A renovated knowledgebase for deriving biological and chemical insight from metabolites, proteins, and genes. Bioinformatics, 39(1):btac726, 2023. ISSN 1367-4811. doi: 10.1093/bioinformatics/btac726.

[5] Minoru Kanehisa, Miho Furumichi, Mao Tanabe, Yoko Sato, and Kanae Morishima. KEGG: New perspectives on genomes, pathways, diseases and drugs. Nucleic Acids Research, 45 (D1):D353–D361, January 2017. ISSN 1362-4962. doi: 10.1093/nar/gkw1092.

[6] Reactome Pathway Knowledgebase 2024 | Nucleic Acids Research | Oxford Academic. https://academic.oup.com/nar/article/52/D1/D672/7369850.

[7] Kil Sun Lee, Xiaoyang Su, and Tao Huan. Metabolites are not genes — avoiding the misuse of pathway analysis in metabolomics. Nature Metabolism, 7(5):858–861, May 2025. ISSN 2522-5812. doi: 10.1038/s42255-025-01283-0.

[8] D Dimitrov, PSL Schäfer, E Farr, P Rodriguez-Mier, S Lobentanzer, P Badia-I-Mompel, A Dugourd, J Tanevski, RO Ramirez Flores, and J Saez-Rodriguez. LIANA+ provides an all-in-one framework for cell-cell communication inference. Nature Cell Biology, 26(9): 1613–1622, 2024. ISSN 1476-4679. doi: 10.1038/s41556-024-01469-w.

[9] E Farr, D Dimitrov, C Schmidt, D Turei, S Lobentanzer, A Dugourd, and J Saez-Rodriguez. MetalinksDB: A flexible and contextualizable resource of metabolite-protein interactions. Briefings in Bioinformatics, 25(4), 2024. doi: 10.1093/bib/bbae347.

[10] Richard W. Carthew. Gene Regulation and Cellular Metabolism: An Essential Partnership. Trends in genetics: TIG, 37(4):389–400, April 2021. ISSN 0168-9525. doi: 10.1016/j.tig.2020.09.018.

[11] Joshua D. Rabinowitz and Thomas J. Silhavy. Systems biology: Metabolite turns master regulator. Nature, 500(7462):283–284, August 2013. ISSN 1476-4687. doi: 10.1038/nature12544.

[12] Christian M. Metallo and Matthew G. Vander Heiden. Understanding metabolic regulation and its influence on cell physiology. Molecular Cell, 49(3):388–398, February 2013. ISSN 1097-4164. doi: 10.1016/j.molcel.2013.01.018.

[13] Juan Manuel Schvartzman, Craig B. Thompson, and Lydia W. S. Finley. Metabolic regulation of chromatin modifications and gene expression. The Journal of Cell Biology, 217(7): 2247–2259, July 2018. ISSN 1540-8140. doi: 10.1083/jcb.201803061.

[14] Identification of bioactive metabolites using activity metabolomics | Nature Reviews Molecular Cell Biology. https://www.nature.com/articles/s41580-019-0108-4.

[15] Dénes Türei, Tamás Korcsmáros, and Julio Saez-Rodriguez. OmniPath: Guidelines and gateway for literature-curated signaling pathway resources. Nature Methods, 13(12):966–967, December 2016. ISSN 1548-7105. doi: 10.1038/nmeth.4077.

[16] Dénes Türei, Jonathan Schaul, Nicolàs Palacio-Escat, Balázs Bohár, Yunfan Bai, Francesco Ceccarelli, Elif Çevrim, Macabe Daley, Melih Darcan, Daniel Dimitrov, Tunca Doğan, Daniel Domingo-Fernández, Aurelien Dugourd, Attila Gábor, Lejla Gul, Benjamin A Hall, Charles Tapley Hoyt, Olga Ivanova, Michal Klein, Toby Lawrence, Diego Mañanes, Dezső Módos, Sophia Müller-Dott, Márton Ölbei, Christina Schmidt, Bünyamin Şen, Fabian J Theis, Atabey Ünlü, Erva Ulusoy, Alberto Valdeolivas, Tamás Korcsmáros, and Julio Saez-Rodriguez. OmniPath: Integrated knowledgebase for multi-omics analysis. Nucleic Acids Research, 54(D1):D652–D660, January 2026. ISSN 1362-4962. doi: 10.1093/nar/gkaf1126.

[17] RDKit: Open-source cheminformatics and Machine Learning.

[18] Adnan Malik, Muhammad Arsalan, Carlos Moreno, Juan Mosquera, Eloy Félix, Tevfik Kizilören, Venkatesh Muthukrishnan, Barbara Zdrazil, Andrew R Leach, and Noel M O’Boyle. ChEBI: Re-engineered for a sustainable future. Nucleic Acids Research, 54(D1): D1768–D1778, November 2025. ISSN 0305-1048. doi: 10.1093/nar/gkaf1271.

[19] Kirill Degtyarenko, Paula de Matos, Marcus Ennis, Janna Hastings, Martin Zbinden, Alan McNaught, Rafael Alcántara, Michael Darsow, Mickaël Guedj, and Michael Ashburner. ChEBI: A database and ontology for chemical entities of biological interest. Nucleic Acids Research, 36(Database issue):D344–350, January 2008. ISSN 1362-4962. doi: 10.1093/nar/gkm791.

[20] Dominik Kopczynski, Nils Hoffmann, Bing Peng, Gerhard Liebisch, Friedrich Spener, and Robert Ahrends. Goslin 2.0 Implements the Recent Lipid Shorthand Nomenclature for MS-Derived Lipid Structures. Analytical Chemistry, 94(16):6097–6101, April 2022. ISSN 0003-2700. doi: 10.1021/acs.analchem.1c05430.

[21] Greg Landrum, Paolo Tosco, Brian Kelley, Ricardo Rodriguez, David Cosgrove, Riccardo Vianello, sriniker, Peter Gedeck, Gareth Jones, Dan Nealschneider, Eisuke Kawashima, NadineSchneider, tadhurst-cdd, Andrew Dalke, Matt Swain, Brian Cole, Samo Turk, Aleksandr Savelev, Niels Maeder, Yakov Pechersky, Alain Vaucher, Maciej Wójcikowski, Rachel Walker, Hussein Faara, Ichiru Take, Vincent F. Scalfani, Daniel Probst, Kazuya Ujihara, Jeremy Monat, and Juuso Lehtivarjo. RDKit: Open-source cheminformatics. Zenodo, May 2026.

[22] Anna Gaulton, Louisa J. Bellis, A. Patricia Bento, Jon Chambers, Mark Davies, Anne Hersey, Yvonne Light, Shaun McGlinchey, David Michalovich, Bissan Al-Lazikani, and John P. Overington. ChEMBL: A large-scale bioactivity database for drug discovery. Nucleic Acids Research, 40(Database issue):D1100–D1107, January 2012. ISSN 0305-1048. doi: 10.1093/nar/gkr777.

[23] Lucila Aimo, Robin Liechti, Nevila Hyka-Nouspikel, Anne Niknejad, Anne Gleizes, Lou Götz, Dmitry Kuznetsov, Fabrice P. A. David, F. Gisou van der Goot, Howard Riezman, Lydie Bougueleret, Ioannis Xenarios, and Alan Bridge. The SwissLipids knowledgebase for lipid biology. Bioinformatics, 31(17):2860–2866, September 2015. ISSN 1367-4811. doi: 10.1093/bioinformatics/btv285.

[24] Michael K. Gilson, Tiqing Liu, Michael Baitaluk, George Nicola, Linda Hwang, and Jenny Chong. BindingDB in 2015: A public database for medicinal chemistry, computational chemistry and systems pharmacology. Nucleic Acids Research, 44(Database issue):D1045–D1053, January 2016. ISSN 0305-1048. doi: 10.1093/nar/gkv1072.

[25] Min-Gyoung Shin and Alexander R. Pico. Using published pathway figures in enrichment analysis and machine learning. BMC Genomics, 24(1):713, November 2023. ISSN 1471-2164. doi: 10.1186/s12864-023-09816-1.

[26] David S Wishart, AnChi Guo, Eponine Oler, Fei Wang, Afia Anjum, Harrison Peters, Raynard Dizon, Zinat Sayeeda, Siyang Tian, Brian L Lee, Mark Berjanskii, Robert Mah, Mai Yamamoto, Juan Jovel, Claudia Torres-Calzada, Mickel Hiebert-Giesbrecht, Vicki W Lui, Dorna Varshavi, Dorsa Varshavi, Dana Allen, David Arndt, Nitya Khetarpal, Aadhavya Sivakumaran, Karxena Harford, Selena Sanford, Kristen Yee, Xuan Cao, Zachary Budinski, Jaanus Liigand, Lun Zhang, Jiamin Zheng, Rupasri Mandal, Naama Karu, Maija Dambrova, Helgi B Schiöth, Russell Greiner, and Vasuk Gautam. HMDB 5.0: The Human Metabolome Database for 2022. Nucleic Acids Research, 50(D1):D622–D631, January 2022. ISSN 0305-1048, 1362-4962. doi: 10.1093/nar/gkab1062.

[27] David S. Wishart, Dan Tzur, Craig Knox, Roman Eisner, An Chi Guo, Nelson Young, Dean Cheng, Kevin Jewell, David Arndt, Summit Sawhney, Chris Fung, Lisa Nikolai, Mike Lewis, Marie-Aude Coutouly, Ian Forsythe, Peter Tang, Savita Shrivastava, Kevin Jeroncic, Paul Stothard, Godwin Amegbey, David Block, David. D. Hau, James Wagner, Jessica Miniaci, Melisa Clements, Mulu Gebremedhin, Natalie Guo, Ying Zhang, Gavin E. Duggan, Glen D. MacInnis, Alim M. Weljie, Reza Dowlatabadi, Fiona Bamforth, Derrick Clive, Russ Greiner, Liang Li, Tom Marrie, Brian D. Sykes, Hans J. Vogel, and Lori Querengesser. HMDB: The Human Metabolome Database. Nucleic Acids Research, 35(suppl_1):D521–D526, January 2007. ISSN 0305-1048. doi: 10.1093/nar/gkl923.

[28] S Kim, PA Thiessen, EE Bolton, J Chen, G Fu, A Gindulyte, L Han, J He, S He, BA Shoemaker, J Wang, B Yu, J Zhang, and SH Bryant. PubChem Substance and Compound databases. Nucleic Acids Research, 44(D1):D1202–13, 2016. doi: 10.1093/nar/gkv951.

[29] Matthew JConroy, Robert M Andrews, Simon Andrews, Lauren Cockayne, Edward A Dennis, Eoin Fahy, Caroline Gaud, William J Griffiths, Geoff Jukes, Maksim Kolchin, Karla Mendivelso, Andrea F Lopez-Clavijo, Caroline Ready, Shankar Subramaniam, and Valerie B O’Donnell. LIPID MAPS: Update to databases and tools for the lipidomics community. Nucleic Acids Research, 52(D1):D1677–D1682, January 2024. ISSN 0305-1048. doi: 10.1093/nar/gkad896.

[30] Manish Sud, Eoin Fahy, Dawn Cotter, Alex Brown, Edward A. Dennis, Christopher K. Glass, Alfred H. Merrill, Robert C. Murphy, Christian R. H. Raetz, David W. Russell, and Shankar Subramaniam. LMSD: LIPID MAPS structure database. Nucleic Acids Research, 35 (Database issue):D527–532, January 2007. ISSN 1362-4962. doi: 10.1093/nar/gkl838.

[31] Eoin Fahy and Shankar Subramaniam. RefMet: A reference nomenclature for metabolomics. Nature Methods, 17(12):1173–1174, December 2020. ISSN 1548-7105. doi: 10.1038/s41592-020-01009-y.

[32] Donna Maglott, Jim Ostell, Kim D. Pruitt, and Tatiana Tatusova. Entrez Gene: Gene-centered information at NCBI. Nucleic Acids Research, 33(Database issue):D54–58, January 2005. ISSN 1362-4962. doi: 10.1093/nar/gki031.

[33] [33] Aurelien Dugourd, Pascal Lafrenz, Diego Mañanes, Victor Paton, Robin Fallegger, Anne-Claire Kroger, Denes Turei, Yunfan Bai, Yuxin Li, Michael Trogdon, Drew Nager, Shib-ing Deng, Chen Shen, John D. Lapek, Blerta Shtylla, and Julio Saez-Rodriguez. Modeling causal signal propagation in multiomic factor space with COSMOS, July 2024.

[34] Aurelien Dugourd, Christoph Kuppe, Marco Sciacovelli, Enio Gjerga, Attila Gabor, Kristina B. Emdal, Vitor Vieira, Dorte B. Bekker-Jensen, Jennifer Kranz, Eric M. J. Bindels, Ana S. H. Costa, Abel Sousa, Pedro Beltrao, Miguel Rocha, Jesper V. Olsen, Christian Frezza, Rafael Kramann, and Julio Saez-Rodriguez. Causalintegrationofmulti-omicsdata with prior knowledge to generate mechanistic hypotheses. Molecular Systems Biology, 17 (1):e9730, January 2021. ISSN 1744-4292. doi: 10.15252/msb.20209730.

[35] D Shorthouse, J Bradley, SE Critchlow, C Bendtsen, and BA Hall. Heterogeneity of the cancer cell line metabolic landscape. Molecular Systems Biology, 18(11):e11006, 2022. doi: 10.15252/msb.202211006.

[36] Yanling Sun, Xinchang Zheng, Guoliang Wang, Yibo Wang, Xiaoning Chen, Jiani Sun, Zhuang Xiong, Sisi Zhang, Tianyi Wang, Zhuojing Fan, Congfan Bu, Yiming Bao, and Wen-ming Zhao. MACdb: A Curated Knowledgebase for Metabolic Associations across Human Cancers. Molecular Cancer Research, 21(7):691–697, July 2023. ISSN 1541-7786, 1557–3125. doi: 10.1158/1541-7786.MCR-22-0909.

[37] A. S. Breathnach, M. Nazzaro-Porro, and S. Passi. AZELAIC ACID. British Journal of Dermatology, 111(1):115–120, July 1984. ISSN 0007-0963. doi: 10.1111/j.1365-2133.1984. tb04025.x.

[38] Antonio Ayala, Mario F. Muñoz, and Sandro Argüelles. Lipid peroxidation: Production, metabolism, and signaling mechanisms of malondialdehyde and 4-hydroxy-2-nonenal. Oxidative Medicine and Cellular Longevity, 2014:360438, 2014. ISSN 1942-0994. doi: 10.1155/2014/360438.

[39] Gina M. DeNicola, Florian A. Karreth, Timothy J. Humpton, Aarthi Gopinathan, Cong Wei, Kristopher Frese, Dipti Mangal, Kenneth H. Yu, Charles J. Yeo, Eric S. Calhoun, Francesca Scrimieri, Jordan M. Winter, Ralph H. Hruban, Christine Iacobuzio-Donahue, Scott E. Kern, Ian A. Blair, and David A. Tuveson. Oncogene-induced Nrf2 transcription promotes ROS detoxification and tumorigenesis. Nature, 475(7354):106–109, July 2011. ISSN 1476-4687. doi: 10.1038/nature10189.

[40] Zachary A. King, Justin Lu, Andreas Dräger, Philip Miller, Stephen Federowicz, Joshua A. Lerman, Ali Ebrahim, Bernhard O. Palsson, and Nathan E. Lewis. BiGG Models: A platform for integrating, standardizing and sharing genome-scale models. Nucleic Acids Research, 44(D1):D515–D522, January 2016. ISSN 0305-1048. doi: 10.1093/nar/gkv1049.

[41] JL Robinson, P Kocabaş, H Wang, PE Cholley, D Cook, A Nilsson, M Anton, R Ferreira, I Domenzain, V Billa, A Limeta, A Hedin, J Gustafsson, EJ Kerkhoven, LT Svensson, BO Palsson, A Mardinoglu, L Hansson, M Uhlén, and J Nielsen. An atlas of human metabolism. Science Signaling, 13(624), 2020. ISSN 1945-0877. doi: 10.1126/scisignal. aaz1482.

[42] Julia Hauenstein, Lisa Jeske, Antje Jäde, Mathias Krull, Katrin Dümmer, Julia Koblitz, Anja Tietz, Dieter Jahn, Lorenz Christian Reimer, and Boyke Bunk. BRENDA in 2026: A Global Core Biodata Resource for functional enzyme and metabolic data within the DSMZ Digital Diversity. Nucleic Acids Research, 54(D1):D527–D534, January 2026. ISSN 0305-1048, 1362-4962. doi: 10.1093/nar/gkaf1113.

[43] Reza M Salek, Christoph Steinbeck, Mark R Viant, Royston Goodacre, and Warwick B Dunn. The role of reporting standards for metabolite annotation and identification in metabolomic studies. GigaScience, 2(1):13, December 2013. ISSN 2047-217X. doi: 10.1186/2047-217X-2-13.

[44] Pablo Rodriguez-Mier, Martin Garrido-Rodriguez, Attila Gabor, and Julio Saez-Rodriguez. Unifying multi-sample network inference from prior knowledge and omics data with CORNETO. Nature Machine Intelligence, 7(7):1168–1186, July 2025. ISSN 2522-5839. doi: 10.1038/s42256-025-01069-9.

[45] A Dugourd, P Lafrenz, D Mañanes, R Fallegger, AC Kroger, D Turei, B Shtylla, and J Saez-Rodriguez. Modeling causal signal propagation in multi-omic factor space with COSMOS. BioRxiv, 2024. doi: 10.1101/2024.07.15.603538.

[46] Samuel Kerrien, Sandra Orchard, Luisa Montecchi-Palazzi, Bruno Aranda, Antony F. Quinn, Nisha Vinod, Gary D. Bader, Ioannis Xenarios, Jérôme Wojcik, David Sherman, Mike Tyers, John J. Salama, Susan Moore, Arnaud Ceol, Andrew Chatr-Aryamontri, Matthias Oesterheld, Volker Stümpflen, Lukasz Salwinski, Jason Nerothin, Ethan Cerami, Michael E. Cusick, Marc Vidal, Michael Gilson, John Armstrong, Peter Woollard, Christopher Hogue, David Eisenberg, Gianni Cesareni, Rolf Apweiler, and Henning Hermjakob. Broadening the horizon–level 2.5 of the HUPO-PSI format for molecular interactions. BMC biology, 5:44, October 2007. ISSN 1741-7007. doi: 10.1186/1741-7007-5-44.

[47] Mark Raasveldt and Hannes Mühleisen. DuckDB: An Embeddable Analytical Database. In Proceedings of the 2019 International Conference on Management of Data, pages 1981–1984, Amsterdam Netherlands, June 2019. ACM. ISBN 978-1-4503-5643-5. doi: 10.1145/3299869.3320212.

[48] The UniProt Consortium. UniProt: The universal protein knowledgebase in 2021. Nucleic Acids Research, 49(D1):D480–D489, January 2021. ISSN 0305-1048. doi: 10.1093/nar/gkaa1100.

[49] Jon Chambers, Mark Davies, Anna Gaulton, Anne Hersey, Sameer Velankar, Robert Petryszak, Janna Hastings, Louisa Bellis, Shaun McGlinchey, and John P. Overington. UniChem: A unified chemical structure cross-referencing and identifier tracking system. Journal of Cheminformatics, 5(1):3, January 2013. ISSN 1758-2946. doi: 10.1186/1758-2946-5-3.

[50] Sébastien Moretti, Van Du T. Tran, Florence Mehl, Mark Ibberson, and Marco Pagni. MetaNetX/MNXref: Unified namespace for metabolites and biochemical reactions in the context of metabolic models. Nucleic Acids Research, 49(D1):D570–D574, January 2021. ISSN 1362-4962. doi: 10.1093/nar/gkaa992.

[51] Milton H. Saier, Jr, Can V. Tran, and Ravi D. Barabote. TCDB: The Transporter Classification Database for membrane transport protein analyses and information. Nucleic Acids Research, 34(suppl_1):D181–D186, January 2006. ISSN 0305-1048. doi: 10.1093/nar/gkj001.

[52] Milton H Saier, Vamsee S Reddy, Gabriel Moreno-Hagelsieb, Kevin J Hendargo, Yichi Zhang, Vasu Iddamsetty, Katie Jing Kay Lam, Nuo Tian, Steven Russum, Jianing Wang, and Arturo Medrano-Soto. The Transporter Classification Database (TCDB): 2021 update. Nucleic Acids Research, 49(D1):D461–D467, November 2020. ISSN 0305-1048. doi: 10.1093/nar/gkaa1004.

[53] Y Zhang, Y Yang, L Ren, M Zhan, T Sun, Q Zou, and Y Zhang. Predicting intercellular communication based on metabolite-related ligand-receptor interactions with MRCLinkdb. BMC Biology, 22(1):152, 2024. doi: 10.1186/s12915-024-01950-w.

[54] SD Harding, JF Armstrong, E Faccenda, C Southan, SPH Alexander, AP Davenport, AJ Pawson, M Spedding, JA Davies, and NC-IUPHAR. The IUPHAR/BPS guide to PHARMACOLOGY in 2022: Curating pharmacology for COVID-19, malaria and antibacterials. Nucleic Acids Research, 50(D1):D1282–D1294, 2022. doi: 10.1093/nar/gkab1010.

[55] Simon D Harding, Jane F Armstrong, Elena Faccenda, Christopher Southan, Stephen P H Alexander, Anthony P Davenport, Michael Spedding, and Jamie A Davies. The IUPHAR/BPS Guide to PHARMACOLOGY in 2026. Nucleic Acids Research, 54(D1): D1446–D1456, October 2025. ISSN 0305-1048. doi: 10.1093/nar/gkaf1067.

[56] D Türei, A Valdeolivas, L Gul, N Palacio-Escat, M Klein, O Ivanova, M Ölbei, A Gábor, F Theis, D Módos, T Korcsmáros, and J Saez-Rodriguez. Integrated intra- and intercellular signaling knowledge for multicellular omics analysis. Molecular Systems Biology, 17 (3):e9923, 2021. ISSN 1744-4292. doi: 10.15252/msb.20209923.

[57] Bingxue Lyu, Ke Wu, Yuanyuan Huang, Mihail Anton, Xiongwen Li, Sandra Viknander, Danish Anwer, Yunfeng Yang, Diannan Lu, Eduard Kerkhoven, Aleksej Zelezniak, Dan Gao, Yu Chen, and Feiran Li. GotEnzymes2: Expanding coverage of enzyme kinetics and thermal properties. Nucleic Acids Research, 54(D1):D583–D592, January 2026. ISSN 0305-1048, 1362-4962. doi: 10.1093/nar/gkaf1053.

[58] D Szklarczyk, A Santos, C von Mering, LJ Jensen, P Bork, and M Kuhn. STITCH 5: Aug-mentingprotein-chemicalinteractionnetworkswithtissueandaffinitydata. Nucleic Acids Research, 44(D1):D380–4, 2016. doi: 10.1093/nar/gkv1277.

[59] Hikaru Sugimoto, Keigo Morita, Dongzi Li, Yunfan Bai, Matthias Mattanovich, and Shinya Kuroda. iTraNet: A web-based platform for integrated trans-omics network visualization and analysis. Bioinformatics Advances, 4(1):vbae141, January 2024. ISSN 2635-0041. doi: 10.1093/bioadv/vbae141.

[60] Structural robustness and temporal vulnerability of the starvation-responsive metabolic network in healthy and obese mouse liver | Science Signaling. https://www.science.org/doi/10.1126/scisignal.ads2547?url_ver=Z39.88-2003&rfr_id=ori:rid:crossref.org&rfr_dat=cr_pub%20%200pubmed.

[61] Adrián César-Razquin, Berend Snijder, Tristan Frappier-Brinton, Ruth Isserlin, Gergely Gyimesi, Xiaoyun Bai, Reinhart A. Reithmeier, David Hepworth, Matthias A. Hediger, Aled M. Edwards, and Giulio Superti-Furga. A Call for Systematic Research on Solute Carriers. Cell, 162(3):478–487, July 2015. ISSN 0092-8674. doi: 10.1016/j.cell.2015.07.022.

[62] Sandra Placzek, Ida Schomburg, Antje Chang, Lisa Jeske, Marcus Ulbrich, Jana Tillack, and Dietmar Schomburg. BRENDA in 2017: New perspectives and new tools in BRENDA. Nucleic Acids Research, 45(D1):D380–D388, January 2017. ISSN 1362-4962. doi: 10.1093/nar/gkw952.

[63] Amos Bairoch. The Cellosaurus, a Cell-Line Knowledge Resource. Journalof Biomolecular Techniques : JBT, 29(2):25–38, July 2018. ISSN 1524-0215. doi: 10.7171/jbt.18-2902-002.

[64] Christina Schmidt, Denes Turei, Dimitrios Prymidis, Macabe Daley, Christian Frezza, and Julio Saez-Rodriguez. Integrated metabolomics data analysis to generate mechanistic hypotheses with MetaProViz, August 2025. ISSN 2692-8205.

[65] Matthew E. Ritchie, Belinda Phipson, Di Wu, Yifang Hu, Charity W. Law, Wei Shi, and Gordon K. Smyth. Limma powers differential expression analyses for RNA-sequencing and microarray studies. Nucleic Acids Research, 43(7):e47, April 2015. ISSN 1362-4962. doi: 10.1093/nar/gkv007.

[66] Yoav Benjamini and Yosef Hochberg. Controlling the False Discovery Rate: A Practical and Powerful Approach to Multiple Testing. Journal of the Royal Statistical Society: Series B (Methodological), 57(1):289–300, 1995. ISSN 2517-6161. doi: 10.1111/j.2517-6161.1995.tb02031.x.

[67] Elizabeth Brunk, Swagatika Sahoo, Daniel C. Zielinski, Ali Altunkaya, Andreas Dräger, Nathan Mih, Francesco Gatto, Avlant Nilsson, German Andres Preciat Gonzalez, Maike Kathrin Aurich, Andreas Prlić, Anand Sastry, Anna D. Danielsdottir, Almut Heinken, Alberto Noronha, Peter W. Rose, Stephen K. Burley, Ronan M. T. Fleming, Jens Nielsen, Ines Thiele, and Bernhard O. Palsson. Recon3D enables a three-dimensional view of gene variation in human metabolism. Nature Biotechnology, 36(3):272–281, March 2018. ISSN 1546-1696. doi: 10.1038/nbt.4072.

